# Development alters genotype-environment interactions and shapes adaptation in *Arabidopsis*

**DOI:** 10.1101/2025.05.13.653704

**Authors:** Erica H Lawrence-Paul, Jessica Janakiraman, Matthew R Lawrence-Paul, Rotem Ben-Zeev, Yuxing Xu, Amanda Penn, Jesse R Lasky

## Abstract

The timing of developmental transitions is critical for adaptation, aligning stress-tolerant and sensitive stages with harsh and benign periods. At the same time, tradeoffs can promote diversity in development within populations. While geographic clines in developmental timing are often documented, there are few documented genetic tradeoffs in stress tolerance across developmental stages. Here, we study the juvenile-to-adult vegetative phase change (VPC), a conserved transition marked by changes in leaf morphology and physiology. In Arabidopsis, we found strong rank-changing genotype-by-environment interactions for fitness depending on the stage of drought exposure, indicating tradeoffs and stage-specific mechanisms of drought adaptation. Adult plants significantly increased water-use efficiency under drought, while juveniles did not, indicating lower juvenile plasticity. VPC timing varied with climate in Iberia, where genotypes from warm, dry climates transitioned earlier than those from cool, moist climates. However, SNPs associated with VPC timing and drought fitness showed little global geographic variation, suggesting tradeoffs maintain diversity across the species range. Genome-wide association mapping revealed candidate loci for VPC timing and stage-specific drought responses, several validated in T-DNA lines. Our results show that VPC timing contributes to drought adaptation and that genetic tradeoffs across developmental stages help maintain natural diversity.

## Introduction

The timing and nature of developmental transitions is critical for organismal adaptation. As organisms transition through developmental phases their morphology and physiology changes, potentially altering the way they respond to environmental stress. Understanding the regulation of developmental timing as well as how distinct developmental phases modulate physiology is essential for predicting how organisms will respond to changing climates and for identifying genes and traits important for stress tolerance across lifespans ^1^.

Developmental transitions are deeply conserved in their underlying genetics and phenotypic shifts, yet paradoxically they also represent a key axis of diversification among species and populations. In plants, flowering time and germination are well known contributors to environmental adaptation ^2–7^. The timing of these developmental transitions dictates which organs are exposed to specific environmental conditions, and particularly the environment experienced when shifting from producing vegetative to reproductive structures. While there is an abundance of evidence that the timing of developmental transitions is important for plant adaptation, **few studies have addressed how developmental changes in phenotypes affect fitness, in particular, across environmental gradients**.

We recently showed that juvenile vegetative *Arabidopsis* plants have reduced phenotypic plasticity in response to multiple abiotic gradients, compared to adult vegetative plants, and there is substantial natural variation in stage-specific plasticity ^1^. However, it remains unknown how selection across environments may affect this diversity, and what evolutionary forces act on the underlying genes. Phase-specific shifts in traits can enable plants to alter ecological strategies across their lifespan. For example, ontogeny-dependent habitat filtering promotes a fast-growing “ruderal” strategy in juveniles, later shifting towards either a competitive or stress-resilient adult strategy in grassland and forest ecosystems respectively ^8^. Phase-specific selection can favor the evolution of traits that are costly in one life stage, but advantageous during another. A striking example comes from divaricate woody plants of New Zealand, which evolved a structurally expensive juvenile branching morphology that deters browsing by ratites. Phase-specific regulation of this defense allows plants to transition to a less costly adult morphology once branches grow above the birds’ reach ^9,10^. Together, these examples highlight how ontogenetic shifts can create opportunities for selection to shape stage-specific strategies.

Phenotypes exhibited at one developmental stage may influence or constrain phenotypes at another stage. Many traits increase fitness at one stage at the expense of fitness at another stage, creating a fitness tradeoff across development ^11^. When these tradeoffs balance, variation in the underlying traits is neutral and this variation will not be eliminated by selection ^12^. Plants are classic systems for studying tradeoffs across development ^13–15^ however, tradeoffs between juvenile and adult vegetative stages, as well as their impact on fitness is not well documented. Vegetative phase change (VPC) the transition from juvenile to adult vegetative stages, is regulated by a highly conserved microRNA, miR156, and its targets, *SQUAMOSA PROMOTER BINDING-LIKE* (*SPL*) transcription factors ^16–18^. In *Arabidopsis,* there are 16 *SPL* genes (10 of which are targets of miR156) that regulate shoot growth and morphology, root development, floral induction, stress responses, ecophysiology, and more ^19–24^. The myriad genes regulated by these ten *SPLs* may allow plants to broadly shift many dimensions of physiology across VPC. The fact that many traits are affected by VPC suggests that different environments could select for different timing of VPC and that selection on the timing of vegetative phase, and associated *SPL*-regulated trait changes, could facilitate local adaptation.

Environmental selection acts not only on the timing of developmental transitions, but also on genes and traits expressed during each developmental phase. Because development controls the expression of traits, it could influence the evolution of mutations controlling those traits. Additionally, we have shown that the capacity for phenotypic plasticity itself changes with developmental stage. Juvenile leaves are more canalized and less plastic in response to abiotic stress compared to adult leaves in *Arabidopsis* ^25^ As a result, adaptation to environmental conditions early in life may involve selection for mutations that affect constitutive trait expression, whereas adaptation to conditions experienced at the adult stage may favor mutations that modify plasticity. This hypothesis may be consistent with the known action of the miR156-SPL regulatory module. miR156 suppresses SPL expression during the juvenile stage, so genes positively regulated by SPLs may experience weak selection at the juvenile-stage. By contrast, these SPLs can act as positive regulators of abiotic stress responses later in development exposing them to potentially variable selection across environments ^20,26,27^. Therefore, mutations affecting stress responsive expression of SPL-regulated genes may only manifest phenotypes in adult plants experiencing those stressors. Overall, we hypothesize that developmental regulation influences which genes are exposed to selection and that while the evolution of expression plasticity likely plays an important role in local adaptation to climate in *Arabidopsis* ^28,29^, genotype-by-environment interactions at the juvenile stage are weaker.

Because few studies have examined plant stress responses across developmental phases -- and even fewer have done so across diverse genotypes -- we know little about how the timing of developmental transitions contributes to local adaptation and tradeoffs through phase-specific responses to abiotic stress. Here we investigate how natural variation in the timing of VPC and phase-specific drought responses contribute to plant adaptation. Using growth chamber experiments with drought treatments applied at different developmental stages in *Arabidopsis thaliana* natural genotypes, along with validation experiments in mutants, we address the following questions:

1. **Are the phenotypic effects of drought specific to each developmental stage?** We hypothesized that juvenile and adult vegetative plants would differ in performance and physiological responses to drought, suggesting developmental shifts in ecological strategy.
2. **How does natural selection act on the timing of development and stage-specific drought?** We hypothesized that geographically variable or balancing selection shapes developmental timing and phase-specific traits so that plants are well suited to tolerate environmental stress during the life stages when those stressors are most likely to occur.
3. **Are there tradeoffs in drought adaptation across developmental stages?** Tradeoffs could maintain diversity in developmental timing between or within populations. We hypothesized that developmental shifts in ecological strategy would cause different traits to underlie drought adaptation in each stage.
4. **What genes underlie genetic variation in the timing of development and stage-specific drought adaptation?** We hypothesized that genes regulated by, or regulating SPL transcription factors and miR156/7 are especially likely to contribute, since expression of these genes likely varies dramatically between juvenile and adult phases.

## Results

### Developmental stage determines rosette growth response to drought

Plants of 158 ecotypes were subjected to a 10-day drought during either the juvenile or adult phases of vegetative development, or were well-watered across their entire lifespan as a control. Rosette growth responses to drought differed between juvenile and adult phases. In the juvenile phase, drought significantly reduced the amount of growth (*p* = 2.2e^-16^) whereas there was no significant impact on growth during the adult phase (*p* = 0.078) (Fig 1a). Plants grew an average of 1.37 cm (±0.62) in diameter less in drought compared to control in the juvenile phase whereas during the adult drought treatment, plants grew nearly the same amount (0.28 ±0.93 cm less) as well-watered controls. Thus, the stage when drought occurs impacts growth in a phase-specific way with juvenile plants being more sensitive to drought than adult plants. Interestingly, there was significant GxE (*p* = 0.001) between growth during drought in the juvenile and adult phases, indicating that drought tolerance is phase-specific as genotypes with high growth rates during drought were not consistent between developmental phases.

**Figure 1.**
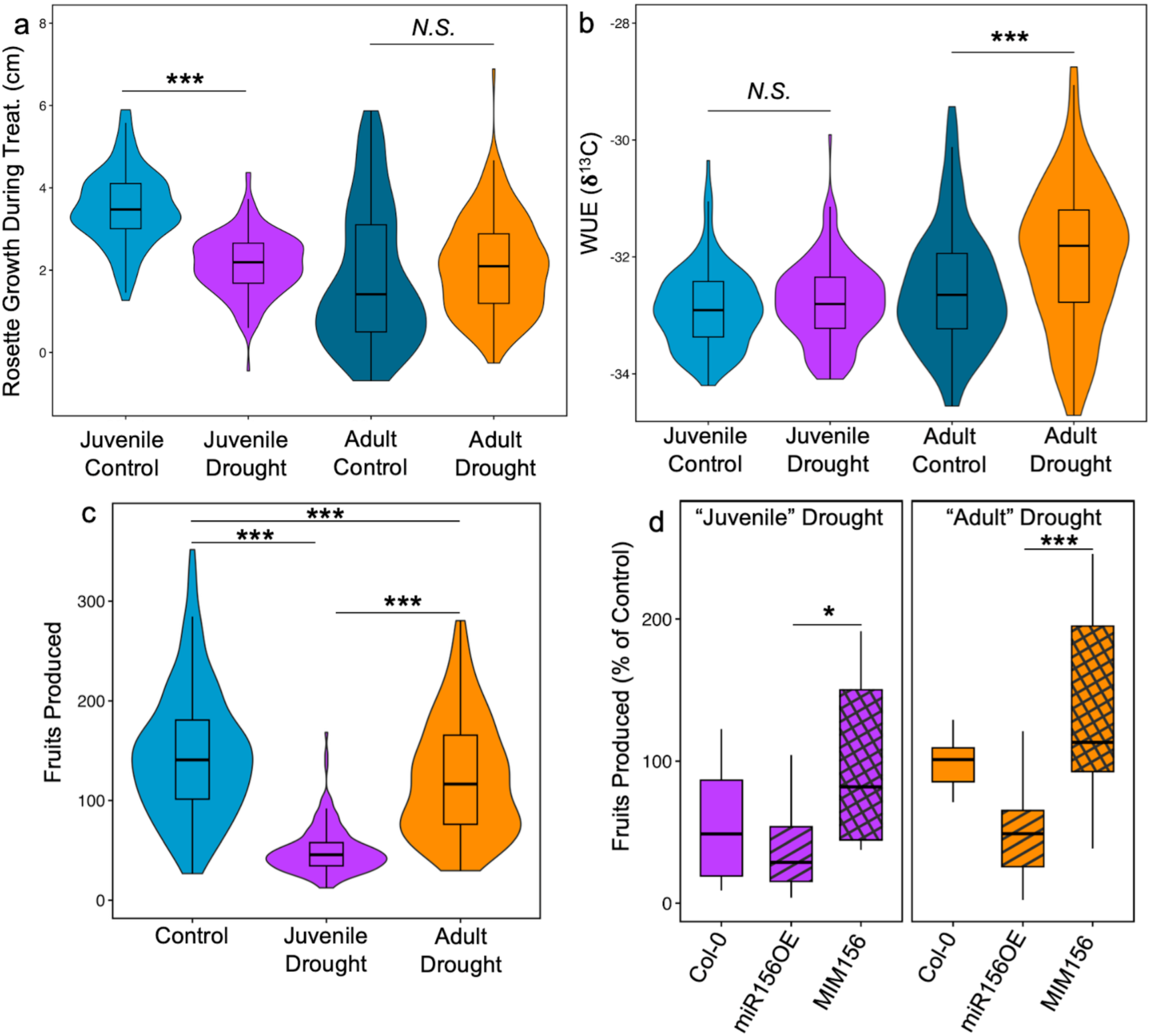
Rosette diameter growth during drought treatments (A), carbon isotope discrimination representing water use efficiency (WUE) (B) and fruits produced (C), shows phase-specific responses to drought. Box plots display the median and interquartile range for traits of plants in well-watered control, juvenile drought treatments and adult drought plants. Violin plots show the distribution of the data for all 158 genotypes with thick sections indicating higher frequency and thinner sections lower frequency values. (D) shows the role of miR156 expression on phase-specific fitness response to drought using Col-0 wildtype, miR156 overexpression (miR156OE) and MIM156 mutants with reduced miR156. Drought treatments are based on when Col-0 is in the juvenile and adult phases, however miR156OE lines are always developmentally juvenile while MIM156 lines are adult. Box plots display the median and interquartile range for the number of fruits produced in either juvenile drought (purple) or adult drought (orange) treatments as a % of fruits produced in the well water control treatment. Results from Student’s t-tests between groups shown above each comparison where ***, **, and * indicate *p <* 0.001, 0.01 and 0.05 respectively.

### Development alters drought effects on water use efficiency (WUE)

Leaves were sampled for carbon isotope analysis at the end of drought treatments for plants exposed to drought during juvenile and adult phases as well as at corresponding timepoints from plants in the well-watered control treatment (referred to as juvenile control and adult control). During drought treatments, soil water content decreased to an average of 32.5% and 28.2% of maximum capacity in juvenile and adult drought respectively whereas well-watered control plants saw a minimum soil water content of 71.8% (Fig. S1). Using carbon isotope composition (δ^13^C) as a proxy for water use efficiency (WUE), we found that drought affected WUE differently depending on the plant’s developmental stage. Despite the large growth reduction under juvenile drought, WUE did not change significantly in response to drought (p = 0.16). By contrast in the adult phase, drought led to a significant increase in WUE (*p* <0.0001; Fig. 1bc). The median δ^13^C increased by only 0.11‰ between well-watered and drought-stressed plants in the juvenile phase, compared to an increase of 0.84‰ in the adult phase.

Genetic variation in WUE also increased in the adult phase. The interquartile range (IQR) of δ^13^C values was 1.29‰ and 1.58‰ for adult plants in the control group and adult drought treatments respectively, compared to 0.95‰ and 0.875‰ in juvenile control plants and juvenile drought plants. This suggests greater plasticity and greater genetic variation in WUE during the adult phase. As with rosette growth during drought, GxE was significant (p = 1.94e^-7^) for WUE during juvenile and adult drought indicating genotypic variation in WUE was dependent on the stage when drought was experienced and that WUE responses to drought were phase-specific.

### Development impacts covariation in trait responses to drought

We conducted a principal component analysis (PCA) on developmental, growth, fitness and physiological traits measured across all accessions under three conditions, juvenile drought, adult drought and well-watered control (Fig. S2). This analysis allowed us to examine how these phenotypes covary across genotypes and treatments. Consistent with previous findings, we observed that accessions with delayed flowering tend to exhibit higher WUE, indicating a trade-off between life history timing and water conservation. A similar relationship between VPC and WUE is also observed. Conversely, accessions with greater rosette growth, especially during the adult phase, tended to have lower WUE. Notably, rosette growth under drought conditions, especially during the adult phase, was positively associated with fitness. In contrast, the relationship between growth and fitness was weaker under well-watered conditions. Additionally, we found that the developmental stage of drought stress modulates the relationship between growth and WUE. While these traits are negatively correlated in well-watered conditions, this opposition is reduced under adult drought, and even more so in juvenile drought, suggesting phase-specific differences in the physiological response to drought stress.

### Tradeoffs in drought adaptation between developmental phases

Drought treatments at both juvenile and adult stages resulted in a significant reduction in total fruit production at senescence compared to well-watered controls (ANOVA treatment effect: juvenile drought vs. control *p* < 0.001, adult drought vs. control *p* < 0.01, Fig. 1c). On average, plants produced fewer fruits when drought occurred during the juvenile phase than during the adult phase (ANOVA environment *p* <0.001, Fig. 2c). However, a significant genotype-by-environment interaction (*p* < 0.001) revealed that this pattern was not consistent across genotypes. Weak correlations between fruits produced in each treatment further support genotypic variation in fitness responses to drought (linear regression for fruits produced in control vs juvenile drought R^2^ = 0.08, control vs adult drought R^2^ = 0.06, juvenile drought vs adult drought R^2^ = 0.1).

**Figure 2.**
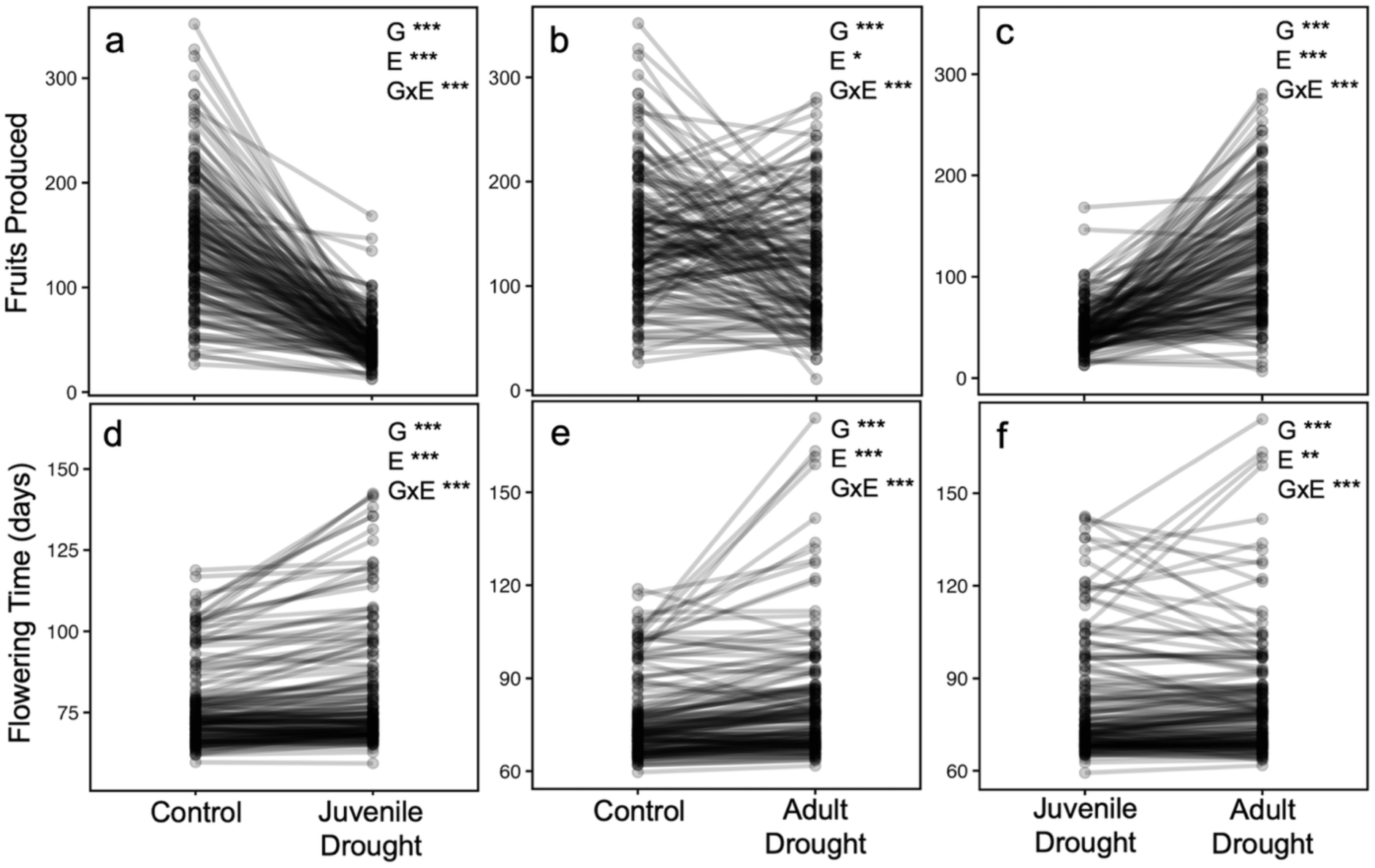
Drought effects on fitness (a-c) and flowering time (d-f) depend on the developmental phase when drought occurs. Black circles indicate the mean number of fruits produced in well-watered control, juvenile drought or adult drought treatments for each genotype and lines connect means between treatments. ANOVA results displayed in the top right corner of each plot where ***, **, and * indicate *p <* 0.001, 0.01 and 0.05 respectively.

For example, accession Aa-0 produced an average of 84 fruits in the juvenile drought treatment but only 30 fruits in adult drought, dropping in fitness rank from 12th to 153rd. In contrast, Ove-0 produced an average of 208 fruits in adult drought but only 43 fruits in juvenile drought, falling from 1st to 90th in rank. Some genotypes such as Krot-0 showed relatively stable performance, producing 147 fruits in juvenile drought and 138 fruits in adult drought. Still, even for such genotypes, the rank order shifted due to more pronounced phase-specific responses in others, causing Krot-0 to drop from a fitness ranking of 2nd in the juvenile drought to 56 in adult drought treatment.

Like with WUE, we saw more genotypic variability in fruit production for plants that experienced drought in the adult phase than in the juvenile phase (IQR juvenile drought = 23.74, adult drought = 89.47). Further supporting the hypothesis that the adult vegetative phase has greater plasticity than the juvenile phase.

These results demonstrate that the developmental timing of drought stress alters its fitness consequences. Shifts in genotype performance and rank order across treatments suggest that the phase in which drought occurs can shape selection and pressures in natural populations.

### Controlling for plant age by using miR156 mutants shows that phase, independent of age, determines fitness responses to drought

miR156OE and MIM156 lines were used to disentangle the effects of plant age from developmental stage as plants overexpressing miR156 remain developmentally juvenile at an age when wild-type plants have transitioned to the adult phase, and plants expressing the MIM156 construct transition to the adult phase when wild-type plants are in the juvenile phase. These lines show that developmental stage, and miR156 expression, independent of plant age, alters drought impacts on fitness. During the juvenile drought treatment, when Col-0 wild-type is in the juvenile phase, adult MIM156 plants with reduced miR156 activity, maintained control levels of fruit production while miR156OE lines with higher levels of miR156 produced an average of only 37.8% of the fruits they were able to produce in well-watered conditions (Tukey adjusted pairwise comparison p = 0.013) (Fig. 1D). In support of this trend, Col-0 plants with natural miR156 levels intermediate to the between the two transgenic lines produced an average of 56.9% of control fruits, although this was not statistically different from the other lines. During the adult drought treatment, when Col-0 wild-type is in the adult phase, MIM156 plants were again able to maintain well-watered fruit production while juvenile miR156OE plants only produced 50.6% of the fruits produced by control plants (Tukey adjusted pairwise comparison p = 0.0011). Again, Col-0 plants showed an intermediate drought response compared to the mutant lines, producing 90% of the control treatment fruit levels, suggesting plants with greater miR156 levels, as is found during the juvenile phase in natural accessions, are more sensitive to drought treatments used in this experiment than those with lower levels of miR156.

### Flowering time plasticity depends on developmental phase during drought

Most genotypes delayed flowering in response to drought during either the juvenile or adult phase, 77% and 79% of genotypes respectively (ANOVA: *p* < 0.001 for both drought treatments, Fig. 2d-e). Under juvenile drought, flowering was delayed by as much as 40.9 days (Tra-01), while some genotypes like Gy-0, flowered as much as 6.6 days earlier than in well-watered control conditions. In the adult drought treatment, shifts in flowering time were even greater for some genotypes with the largest delay of 55.8 days (Sanna-2) and the greatest acceleration of 15.9 days (Ven-1). Significant genotype and genotype-by-environment effects were detected between all treatments (ANOVA G: *p* < 0.001, GxE: *p* < 0.001).

Flowering time responses to drought varied depending on the developmental phase during which it occurred. For example, Sanna-2 showed little change in flowering time under juvenile drought (3.8-day delay) but a strong delay under adult drought (55.8-day delay), resulting in a 52-day difference in flowering time between drought treatments. In contrast, Ost-0 exhibited minimal response under adult drought (0.5-day delay), but a substantial delay under juvenile drought (37.4 days), leading to a 37-day flowering time difference. The significant GxE interaction across all genotypes (ANOVA GxE: *p* <0.001, Fig. 2f) indicates that both the occurrence of drought and the developmental phase during which it is experienced play critical roles in determining flowering time. Because flowering time strongly influences fitness and adaptation, our results suggest that the vegetative stage at which plants encounter abiotic stress may further shape adaptation by altering subsequent developmental transitions.

### WUE associations with fitness and developmental timing are phase-specific

We observed significant but weak negative correlations between WUE and fruit production under drought, indicating that genotypes with higher WUE tended to produce fewer fruits (juvenile drought *p* =0.02, R^2^=0.035; adult drought *p* =0.0018, R^2^ = 0.062). This trade-off may reflect a cost of increased WUE to carbon assimilation, potentially adaptive if drought were extreme enough to cause mortality of low WUE genotypes.

There were also weak but significant correlations between WUE and number of juvenile leaves across several conditions. Genotypes that transitioned earlier to the adult phase tended to have lower WUE (juvenile control: *p* = 0.03, R^2^ = 0.031; adult control: *p* < 0.001, R^2^ = 0.07; adult drought: *p* < 0.001, R^2^ = 0.082). However, this relationship was weaker in well-watered juvenile plants and completely absent under juvenile drought stress (*p* = 0.13, R^2^ = 0.015).

As observed in other studies, higher WUE was associated with later flowering in all growing conditions, reflecting genetic variation in drought avoidance vs. escape strategies (juvenile drought: *p* =2.3e^-5^, R^2^ = 0.11; adult drought *p* = 3.5e^-9^, R^2^ = 0.21; control FT with juvenile control WUE *p* = 1.2e^-5^ R^2^ = 0.12 and control FT with adult control WUE *p* =1.4e^-11^, R^2^= 0.26). Interestingly, flowering time-WUE correlations in the adult phase were stronger than the correlations in the juvenile phase, suggesting this well-known tradeoff is mainly manifested at the adult vegetative stage.

Overall, these results show that developmental stage mediates both the magnitude and variability of WUE responses to drought, with greater plasticity and stronger associations with other traits observed during the adult phase.

### Substantial natural variation in the timing of vegetative phase change

We observed substantial variation in the timing of vegetative phase change (VPC) across the 158 *Arabidopsis* accessions, with transitions as early as leaf 3 or day 48 and as late as leaf 19 or day 64 (Figs. 3, S2). Interestingly, there were no clear geographic patterns in the distribution of VPC timing, suggesting that this diversity is maintained within regions (see analyses below). Two measures of VPC timing, the first leaf with abaxial trichomes, and the number of days from planting until that leaf was visible, were strongly correlated (*p* = 1.6e^-25^, R^2^ = 0.51; Fig. S3b) suggesting little genotypic variation in leaf initiation rate during the juvenile phase. The two morphological indicators of VPC, abaxial trichomes and leaf serrations, were also significantly correlated (*p* = 1.2e^-10^, R^2^ = 0.24; Fig S3c), although some independent variation was evident. This is consistent with previous work that shows the onset of adult traits is staggered due to differential regulation by SPL transcription factors, and varying sensitivity of different SPL genes to miR156 levels ^20,30^.

**Figure 3.**
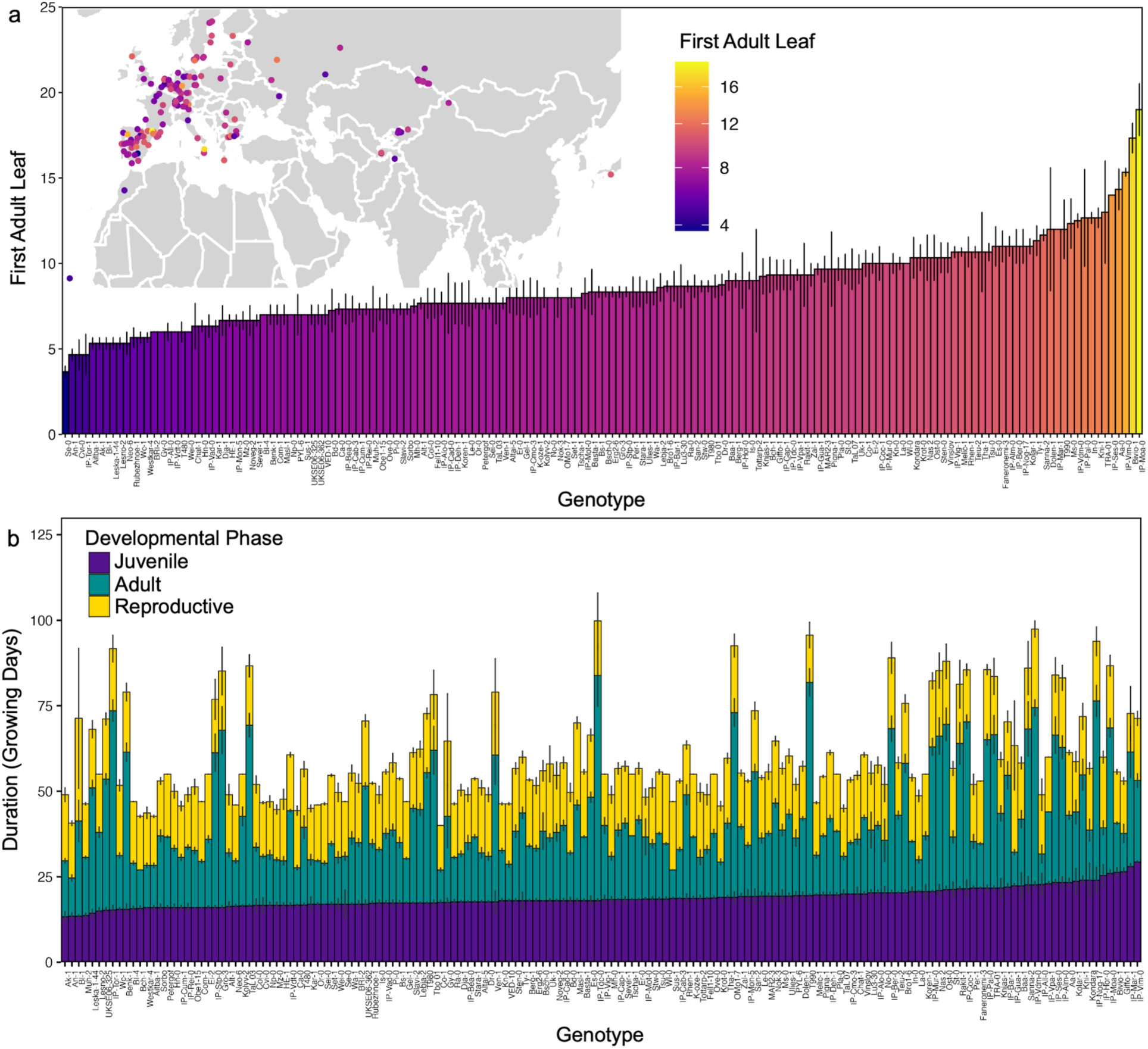
Natural variation in the timing of developmental transitions with genotypes ordered by the timing of vegetative phase change. (A) Timing of vegetative phase change (VPC) based on the first adult leaf with abaxial trichomes for 158 natural accessions used in this study. Accessions are plotted on the map in the location where they originated from, and points are colored based on first adult leaf. Bars represent the mean ± SE for a minimum of 3 replicates per genotype. (B) Duration of time in juvenile vegetative, adult vegetative, and reproductive phases. Bars represent mean ± SE for the duration in days spent in each phase. Juvenile phase, in dark purple, is defined as the time between planting and appearance of first adult leaf (excluding the 35 days spent at 4℃ for vernalization), adult phase (teal) is defined as the time between the appearance of the first adult leaf and first open flower, and reproductive (yellow) is defined as the time between first open flower and first mature fruit

### Independent timing of developmental phases allows flexible life histories

We found that the duration of each developmental phase – juvenile vegetative, adult vegetative, and reproductive (time between flower opening and fruit maturation) – was not strongly correlated (juvenile vs adult: *p* = 0.042, R^2^ = 0.027; juvenile vs reproductive: *p* = 0.98, R^2^ = 4.8e^-6^; adult vs reproductive: *p* = 0.34, R^2^ = 0.0061; Figs. 3b). This indicates that genotypes that transition to the adult phase earlier may compensate by spending more time in the adult vegetative phase before flowering, ultimately reaching reproductive maturity on a similar timescale as genotypes with a slower juvenile transition. For example, Ak-1 spends an average of 13.3 days in the juvenile phase, 16.3 days in the adult phase, and 19.3 days in the reproductive phase for a total of 48.9 days from planting to fruit maturation. In contrast, Giffo-0 spends 26.5 days in the juvenile phase, but has a shorter 11.2 day adult phase and 15.3 day reproductive phase, totaling 53 days. Despite different developmental trajectories, both genotypes complete their life cycle in a similar timeframe suggesting selection on the proportion of time spent in different developmental phases could contribute to local adaptation independent of impacts on total lifecycle length.

### The timing of VPC is stable in response to drought

The timing of VPC showed limited plasticity in response to drought. When abaxial trichomes were used as a marker of VPC, some genotypes showed accelerated transition while others were delayed, however the genotype-by-environment (GxE) interaction was not significant (ANOVA genotype x treatment interaction *p* = 0.47; Fig S3A). On average, plants transitioned at a similar leaf number in both treatments as the average first adult leaf was 8.79 in control and leaf 8.97 in the juvenile drought treatment (ANOVA environment *p* = 0.49). Using leaf serrations as a marker led to similar results for GxE (ANOVA GxE *p* = 0.16; Fig. S4B). However, the overall shift in VPC timing assessed using serrations was slightly delayed under drought, with a mean transition at leaf 7.19 in control and 7.64 in juvenile drought, and this difference was statistically significant (*p* = 0.02). Overall, the limited plasticity in the timing of VPC in response to our drought treatment is reflected in strong correlations between control and drought treatments for both markers (linear regression: trichomes *p* = 6e-05, R^2^ = 0.46; serrations *p* =0.001, R^2^ = 0.33, respectively; Fig S3C-D).

### The timing of VPC is associated with local climate

We analyzed relationships between the timing of vegetative phase change and climate-of-origin, focusing on temperature, precipitation and vapor pressure deficit, during fall (September, October, November) and spring (February, March, April), the main germination and growth seasons for *A. thaliana.* Across all accessions globally, VPC showed no significant association with mean temperature or vapor pressure deficit (VPD) in either season (linear regression between first adult leaf and fall temp: *p =* 0.73, fall VPD: *p* = 0.53, spring temp: *p =* 0.99, spring VPD: *p* = 0.36; Fig4A-B). However, we previously showed that life history clines in Arabidopsis can reverse in direction comparing populations found across the vast native range of Arabidopsis ^31^.

Thus, we also tested for region-specific clines in the timing of VPC. In contrast to global patterns, stronger associations emerged at regional scales. A flexible regression model found a switch from early VPC in warm springs in western Europe to early VPC in cooler spring climates in central Asia (generalized additive model with spatially varying coefficients, Fig. S5). We focused on well sampled regions in Western Europe and found that for accessions from Germany and the Iberian Peninsula, VPC was significantly correlated with both spring and fall temperature, where accessions from warmer areas transitioned to the adult phase earlier than those from cooler regions (Fig 4c,e; Iberia fall mean temp: *p* = 0.007, R^2^ = 0.19; Germany fall mean temp: *p* = 0.02, R^2^ = 0.17; Iberia spring mean temp: *p* = 0.006, R^2^ = 0.2; Germany spring mean temp: *p* = 0.029, R^2^ = 0.16).

**Figure 4.**
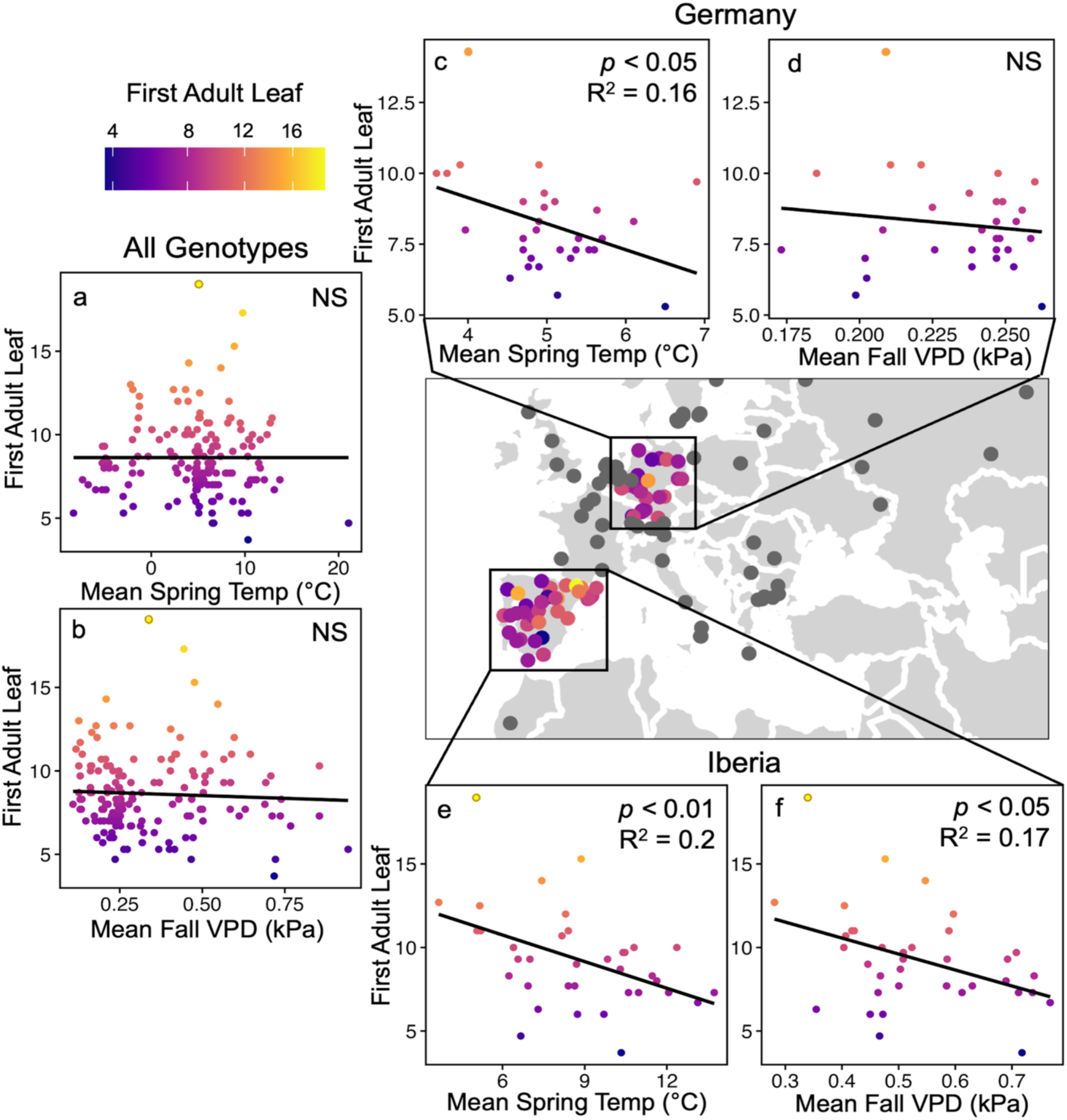
The timing of vegetative phase change is associated with spring temperature and fall vapor pressure deficit (VPD) in native climates. Relationships between the timing of vegetative phase change measured as first leaf with abaxial trichomes and mean spring (March-May) temperature (a, c, e) or mean fall (September-November) VPD (b, d, f) in home environment for all 158 accessions of *Arabidopsis thaliana* used in this study (a, b), accessions from Germany (c, d) or the Iberian peninsula (e, f). Points colored by first adult leaf represent the average per genotype, black lines represent the least-squares regression and results from linear regression analysis are displayed in the top right corner of each plot.

VPD also showed regional associations. In Iberia, accessions from drier areas (higher VPD) transitioned earlier (Fall VPD: *p* = 0.013, R^2^ = 0.165, Spring VPD: *p* = 0.025, R^2^ = 0.14; Fig. 4F). No such relationships were found in Germany (Fall VPD: *p* = 0.5, Spring VPD: *p* = 0.4; Fig 4D), likely because German sites span a much narrower and low VPD range (fall: 0.17-0.26 kPa, spring: 0.14-0.25 kPa) compared with Iberian sites (fall: 0.28-0.77 kPa, spring: 0.17-0.46 kPa). These contrasts suggest that regional climatic heterogeneity can drive local selection on developmental timing, even when global relationships are not significant.

Additional patterns emerged in extreme climates. Accessions from colder regions (i.e. where spring (February, March, April) temperatures average < 0°C, mainly from Asia and Sweden) showed the opposite pattern from Iberia and Germany, warmer origins were associated with later VPC (*p* < 0.01, R^2^ = 0.26). Globally, fall (September, October, November) precipitation was negatively associated with later VPC (*p* = 0.007*, R^2^* = 0.044), with accessions from drier regions transitioning earlier to the more drought-tolerant adult phase. No global relationship was found with spring precipitation (*p* = 0.083). However, in accessions from particularly arid regions (spring precipitations < 100 mm), VPC was strongly correlated with precipitation (*p <* 0.01*, R^2^* = 0.304), with earlier transitions in drier sites. This pattern suggests natural selection may favor earlier VPC under water limitation, aligning the drought-tolerant adult phase with periods of greater stress.

Finally, when we accounted for genome-wide similarity among accessions (using SNPRelate and Kinship2 R packages) these regional significant climate-VPC associations were maintained (spring mean temp Iberia: p = 7e^-5^, Germany: p = 1.8e^-2^, fall mean temp Iberia: : p = 7.8e^-5^, Germany: p = 1.2e^-2^, spring VPD Iberia: p = 4e^-4^, Germany: p = 0.37, fall VPD Iberia: p = 8.1e^-5^, Germany: p = 0.48). This indicates the associations are stronger than expected based on genome-wide evolutionary patterns (which may largely reflect drift), suggesting locally adapted VPC timing traits.

### GWAS identifies novel candidate genes for phase-specific regulators of drought response

We conducted separate genome-wide association studies (GWAS) to identify candidate genes associated with drought responses during the juvenile and adult phases (Fig. 6a-d). While some traits had no single nucleotide polymorphisms (SNPs) surpassing the Storey’s q-value significance threshold, we examined the top SNPs for each trait, recognizing these as potentially informative loci.

To characterize drought tolerance, we used both the total number of fruits produced under drought and the log_2_-fold change in fruits number relative to well-watered controls.

Distinct gene sets were associated with drought tolerance depending on whether stress occurred during the juvenile or adult phase, indicating phase-specific genetic architecture of drought response. Candidate genes near top SNPs in the juvenile phase included *PRA1* (AT3G13720), *Phox/Bem1p* (AT3G18230), *KCS3* (AT1G07720), and *PIP5K8* (AT1G60270), all previously implicated in abiotic stress responses. In the adult phase, candidates included *FBS3* (AT4G05010), *PRLIP7* (AT5G24190), *PRLIP2* (AT5G24200) and *ERF/AP2* (AT2G40350), which have documented roles in drought or stress signaling. Additional candidates included *ARA1* (AT2G40350), involved in starch biosynthesis, and *VPS28-2* (AT4G05000), involved in stomatal closure. We also considered *ERDL4* (AT1G19450), identified in an earlier iteration of our GWAS analysis, and *NCED2* (AT4G18350) which was identified from GWAS for the VPC timing and is a key enzyme in ABA biosynthesis.

To test the roles of these candidates, we analyzed T-DNA insertion lines under juvenile or adult drought treatments, alongside well-watered controls. Five lines (mutants of *PRLIP7, ERF/AP2, NCED2, ARA1, VPS28-2*) showed juvenile-specific effects as mutants produced significantly more fruits than wild-type Col-0 under juvenile drought, but not under adult drought. A mutation in *ERDL4* produced the opposite pattern, conferring fitness advantages under adult drought only. Mutants of PRA1 and Phox/Bem1p (two independent T-DNA lines each) affected drought fitness in both phases, however, Phox/Bem1p lines, showed stronger advantages in the juvenile phase (Fig 5e).

**Figure 5.**
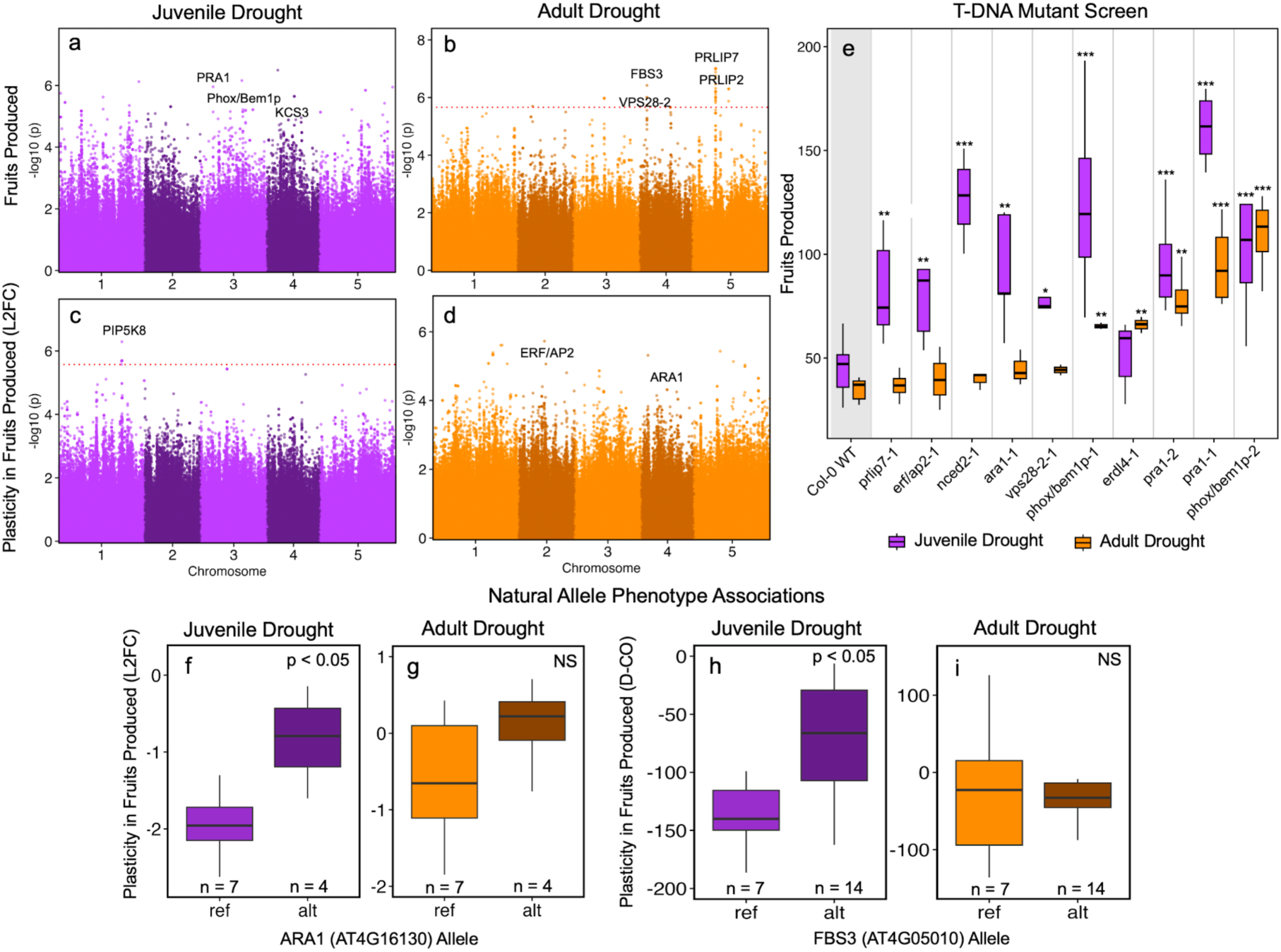
Genetic associations with phase-specific fitness responses. Manhattan plots of genome wide association (GWA) analysis on fitness responses to drought during juvenile and adult phases of development. Manhattan plots show the association of each SNP with the number of fruits produced (a, b) or log2 fold-change in fruits produced between well-watered control and drought treatments (c, d). Red dotted lines indicate significance threshold based on q-values with Benjamin Hochberg FDR correction with *p* < 0.1. Annotations indicate genes associated with top SNPs used in further analyses. (e) Fruits produced by Col-0 wild-type and T-DNA insertion lines when exposed to drought during the juvenile (purple) or adult (orange) phase. Results from Dunnett-adjusted contrasts between Col-0 and each T-DNA line in each condition indicated above each boxplot where ***, **, and * indicate *p <* 0.001, 0.01 and 0.05 respectively. Plasticity in fruit production between well-watered control and juvenile drought (purple) or adult drought (purple) conditions for accessions with the Col-0 reference (ref) allele or predicted loss of function (alt) allele for ARA1 (f-g) or FBS3 (h-i) genes. Plasticity is measured as either the log2 fold-change in fruits produced (f-g) or the difference in fruits produced (h-i). Box plots display the median and interquartile range and number of accessions with each allele denoted at the bottom of each graph. Results from Student’s t-tests between groups shown above each comparison.

To assess whether natural loss of function (LOF) alleles show similar effects, we examined accessions in our study carrying predicted LOF variants. For *ARA1* and *FBS3*, accessions with LOF alleles produced more fruits than reference allele accessions under juvenile drought but performed similarly under adult drought (Fig. 8f-i). This supports a phase-specific benefit of LOF alleles, consistent with our T-DNA results, though *FBS3* T-DNA lines did not significantly differ from Col-0 (Fig. S6a).

Beyond fitness, we identified unique SNPs associated with water-use efficiency (WUE) depending on drought timing (Fig. S7a-b). Juvenile-phase candidates included *RTNLB3* (AT1G64090), linked to plasmodesmata development and *TRM15* (AT4G00440) which is associated with drought response and leaf morphology. Adult-phase candidates included *CCRL9* (AT2G23910), *PPR-like* (AT5G37570) and *PRT* (AT4G36010), all with roles in light responses, as well as *MCA1* (AT4G35920), involved in Ca^2+^ signaling and *CPC* (AT4G35920), involved in epidermal patterning.

Lastly, GWAS for phase-specific flowering time plasticity (Fig. S7c-d) identified multiple candidates, including known effectors of flowering time and fertility (*AUX1, TCP8, AGL13, VIM3*) and genes involved in growth and development (*LACS2, LRR kinase, RDO5*). Together, these results demonstrate that district genetic pathways contribute to drought response at juvenile and adult stages, with different alleles potentially conferring fitness advantages depending on developmental phase.

### GWAS identifies novel candidate genes for the timing of VPC

We conducted a genome-wide association study (GWAS) to identify candidate genes associated with natural variation in the timing of VPC. For the timing of VPC, measured as the first leaf with abaxial trichomes, we identified candidate genes nearest the top-associated SNPs (Fig. 6A). Notably, *CYP71B14* (AT5G25180) is associated with the miR156-SPL regulatory pathway, previously described as a target of *AGL15*, which regulates miR156 expression. Additional candidate genes are involved in cell development, gene silencing, microRNA biogenesis, and DNA and RNA biosynthesis such as *SMR8* (AT1G10690) and *PLAC8* (AT5G05350), *CLASSY3* (AT1G05490), *PRL2* (AT3G05350), and *CTPS4* (AT4G20320) respectively. *NCED2* (AT4G18350) is involved in ABA biosynthesis and has been previously described for having developmental phenotypes. Interestingly, *ROA1* (AT1G60200) which is involved in ABA signaling was identified because of the round juvenile leaf morphology observed in T-DNA insertion line SALK_064472, which is annotated to have a T-DNA insertion in the *PUB19* (AT1G60190) gene, which is just upstream of *ROA1*, that was identified in an early GWAS study for drought response. Zhan *et al.* 2015^32^ Identified that the T-DNA insertion in this line was in fact in *ROA1* and not *PUB19*.

**Figure 6.**
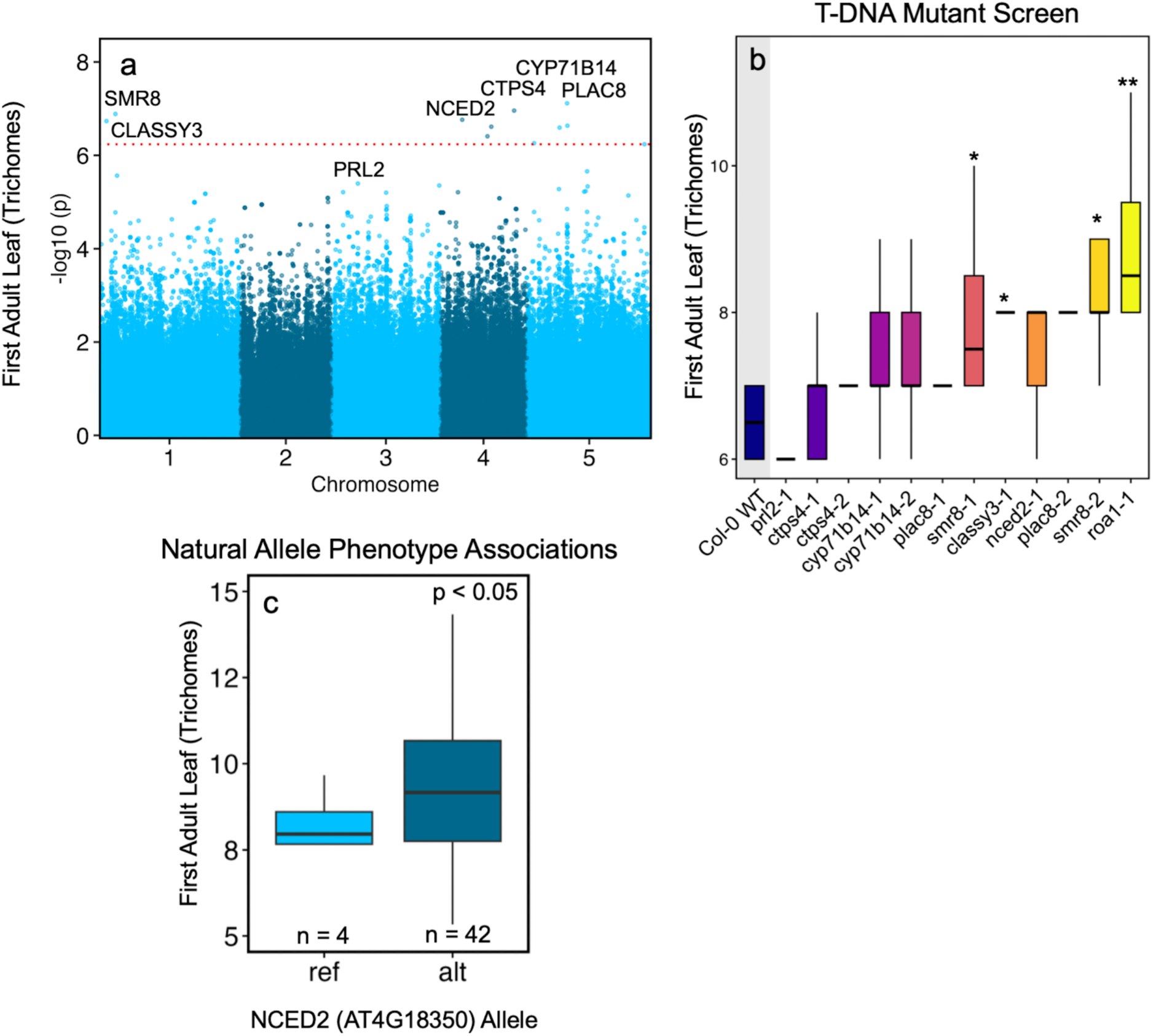
Manhattan plots of genome wide association analysis on the timing of vegetative phase change (VPC) in well-watered control conditions determined as the first leaf with abaxian trichomes. Manhattan plots show the association of each SNP with the node of the first adult leaf. Annotations indicate genes associated with top SNPs used in further analysis. Red dotted lines indicate significance threshold based on q-values with Benjamin Hochberg FDR correction with *p* < 0.1. (b) Position of first adult leaf in Col-0 wild-type and T-DNA insertion lines grown in well-watered control conditions. Results from Dunnett-adjusted contrasts between Col-0 and each T-DNA line indicated above each boxplot where ***, **, and * indicate *p <* 0.001, 0.01 and 0.05 respectively. (c) Position of first adult leaf for accessions with the Col-0 reference (ref) allele or predicted loss of function (alt) allele for NCED2. Box plots display the median and interquartile range and number of accessions with each allele denoted at the bottom of each graph. Results from Student’s t-tests between groups shown at the top of the graph.

We tested the role of these genes using T-DNA insertion lines and measured the first adult leaf with abaxial trichomes. We found that insertions in *SMR8, CLASSY3*, and *ROA1* all lead to significant delays in the onset of adult leaf production (Fig. 6b). As with our drought candidate genes, we looked for accessions in our study with predicted natural loss of function alleles of our genes. *NCED2* had at least 4 accessions included in our study with either the Col-0 reference allele or a natural lof allele. Accessions with the loss of function “alt” allele has significantly delayed VPC (Fig. 6c), and although not statistically significant, the T-DNA insertion line for *NCED2* also showed delayed VPC. Interestingly, our results show that *NCED2* may be involved both in regulating the timing of VPC as well as phase-specific responses to drought.

### Evidence of stage-specific expression and natural selection on GWAS SNPs

Developmental changes in gene expression and silencing may influence the potential for natural variation at such genes to influence stage-specific traits. To test this hypothesis, we used RNA-seq data^33^ to define sets of juvenile- and adult-upregulated genes and examined whether drought-related GWAS SNPs were preferentially located within either set (Fig. S8). Across datasets, GWAS signals for fruit number under both juvenile and adult drought were enriched in genes upregulated during the adult phase, particularly for SNPs with higher minor allele frequencies (MAF=0.4-0.5; adult z=1.98, p=0.001; juvenile z=1.96, p=0.014), suggesting that adaptive drought responses are predominantly associated with genes active during the adult stage. Among GWAS of relative fruit number (log₂ fold change between drought and control), only the adult dataset showed significant enrichment in adult-upregulated genes (all SNPs, z=2.01, p=0.007), and this enrichment was the strongest overall, suggesting that genetic variants underlying drought-response plasticity during the adult phase have greater potential to be targets of natural selection, whereas juvenile responses showed no significant enrichment across any MAF bin (all p>0.09) and potentially likely experience weaker selection.

Spatially varying selection acting on the timing of VPC or stage-specific drought adaptation could result in geographic clines in allele frequency at QTL. We used complementary approaches to characterize geographic variation: the scale-specific genetic variance test, based on wavelet transformed allele frequencies ^34^, and genotype-environment association with climate. First, we found that the top SNPs for the timing of VPC and fruit number under juvenile drought had much less mean geographic variation than expected across a range of spatial scales. Specifically, the timing of VPC top 1000 GWAS SNPs had significantly low spatial variation at the largest scale tested, 2980 km (two tailed permutation test, p<0.02). Additionally, the top 1000 and top 100 SNPs for juvenile fruit number under drought had significantly low spatial variation at this scale (p=0.04 in both cases), and the top 1000 SNPs had low spatial variation at the ∼1360 and ∼620 km scales (p <0.02 and p= 0.04, respectively). These results stand in contrast to our previous findings with flowering time GWAS SNPs, which have highly elevated spatial variation from ∼200-1360 km scales. Fruit number under adult drought SNPs had higher than expected, but not significantly so, spatial variation across scales (all p>0.05).

By contrast, VPC SNPS showed climate GEA enrichments, particularly for precipitation and aridity-related variables. Enrichment was quantified using the Jaccard overlap between top GWAS windows from VPC and climate GEA signals, with 1000 permutation tests to obtain empirical p-values. For fruit number, GWAS signals under both juvenile and adult drought significantly overlapped envGWAS results for water availability, including annual precipitation (adult p=0.041; juvenile p=0.029), precipitation of the driest quarter (adult p=0.017; juvenile p=0.005), and the growing-season aridity index (adult p=0.001; juvenile p=0.003). For plasticity, significant overlap was observed only in the adult dataset for annual precipitation (p=0.001) and precipitation of the driest quarter (p=0.016), whereas the juvenile plasticity GWAS showed no enrichment. These patterns suggest that while water availability shapes the genetic basis of drought performance in both stages, selection on drought-response plasticity is concentrated in the adult phase, whereas juvenile plasticity appears weaker and less influenced by climate-driven selection.

Together, the wavelet and GEA results, though contrasting and using different methods, suggest multiple types of selection acting on genes associated with VPC and stage-specific drought adaptation. These findings that genetic variants associated with traits early in development show less geographic differentiation than expected (i.e. they tend to be similar across populations) suggest they may be subject to balancing selection, whereas variants associated with traits expressed later show more geographic variation indicating selection may act more strongly or more locally on adult-stage traits. Overall, this supports the idea that developmental timing influences how and when natural selection acts on genetic variants.

## Discussion

The developmental stage at which a plant experiences stress plays a critical role in determining how it responds to that stress ^1^. Our results demonstrate that *when* a plant experiences stress can impact fitness as much as *if* it experiences that stress. Developmental phases represent distinct physiological and regulatory states that alter stress resilience, as we found that drought tolerance during the juvenile phase cannot predict tolerance during the adult phase. We present evidence that unique genetic and physiological mechanisms underlie stress resilience across developmental stages, highlighting the importance of studying these phases independently.

The majority of genotypes in our study had lower drought tolerance during the juvenile vegetative phase. Using miR156 mutants, we showed that the miR156-SPL module that regulates vegetative phase change is a driver of these drought response shifts, and that these shifts are developmental phase specific rather than regulated by plant size or age. Since miR156 is highly conserved across the plant kingdom ^35^, findings indicate other economically important crop species will show similar phase-specific drought tolerance

Further, we observed strong genotype-by-environment interactions, with some genotypes more drought-tolerant in the juvenile phase and others more tolerant in the adult phase. This phase-specific GxE suggests that natural selection can act independently on stress responses at different stages. This GxE and the lack of strong negative correlation between performance under drought at the two different stages (linear regression between fruits produced in juvenile vs adult drought R^2^ = 0.1) suggests an opportunity for crop improvement, by selecting for both juvenile and adult phase-specific drought tolerance traits. In this way, breeders could improve stress resilience across the whole lifespan of a plant. While plant biologists already consider pre and post reproductive traits, our results argue that the two vegetative phases warrant similar attention ^36–38^.

We observed considerable natural variation in the timing of development among the accessions used in this study. Notably, the duration of each phase; juvenile, adult and reproductive, varied independently, supporting previous findings that VPC and flowering are independently regulated ^39^. Independent regulation of developmental transitions can allow for developmental phases to align with specific seasonal stress patterns, potentially improving local adaptation. Just as variation in germination and flowering time underlie life history diversity ^4–6^ variation in the timing of VPC may also contribute to different strategies. Previous work links the juvenile phase with fast growth traits like rapid leaf initiation rates, and thin, low-cost tissues, while the adult phase is associated with more expensive tissues with longer lifespans ^24^. Therefore, genotypes that remain in the juvenile phase longer may pursue a “fast” life history strategy focused on growth while those that transition earlier to the adult phase may deploy a strategy focused more on survival.

Interestingly, we observed reduced genetic variation in drought responses during the juvenile phase, consistent with our previous findings that plasticity is reduced early in development compared to the adult phase. One possible explanation is that juvenile traits may be subject to stronger environmental filtering, as this developmental window is closely tied to germination timing, which itself depends on specific environmental cues. To explore this idea, we looked at environmental data from natural *A. thaliana* populations described in Wilczek et al. 2010 ^40^ (Fig. S9). Supporting our hypothesis, we found that temperature and precipitation conditions during early vegetative development more closely resembled those during germination than during reproduction. This suggests that selection may favor more robust, canalized phenotypes during the juvenile phase, reducing plasticity in response to drought and other environmental stresses.

We found the timing of VPC was correlated with climate in Iberia and Germany, suggesting it plays a role in local adaptation, but otherwise we found evidence that selection may maintain diversity in VPC within most populations of *Arabidopsis*. Specifically, there were very weak correlations between the timing of VPC and global climate variation, and the timing of VPC and drought fruit production GWAS SNPs tended to show very little spatial change in allele frequency ^34^. Thus, tradeoffs associated with variation in the timing of VPC, potentially due to variation in abiotic stress response during the juvenile phase, could reduce fitness effects of VPC and promote diversity locally ^12^.

Together, our findings highlight the critical role of developmental transitions, and specifically vegetative phase change, in shaping stress responses and fitness outcomes. A deeper understanding of ontogenetic shifts in morphology and physiology can help to better understand patterns of local adaptation as well as guide the development of more resilient crops to novel climates.

## Methods

### Plant Material

Seeds of 158 *Arabidopsis thaliana* natural accessions (Table S1) were obtained from the *Arabidopsis* Biological Resource Center and grown for seed bulking in a Conviron walk-in growth chamber with 16-hour days at 22°C and 8-hour nights at 18°C with light provided by Philips 32 Watt, 6500K and 4100K fluorescent bulbs and Euri A21 2700K LED bulbs with an average PAR of 194 µmol m^-2^ s^-1^, and 65% relative humidity. Seeds were stratified in 800 ppm Gibberellic acid at 4°C for 5 days then planted in 2 ¼ inch pots filled with Pro-Mix BX growing mix. Pots were covered with humidity domes until after germination and pots were bottom watered regularly to maintain moisture. The natural accessions used were chosen as a subset from those used in Exposito-Alonso *et* al. 2018, where accessions were selected from the 1001 genomes project based on sequence quality and geographic distribution. 35S:miR156a (miR156OE) and 35S:MIM156 lines were obtained from the Poethig Lab at the University of Pennsylvania and have been previously described ^16,41^. T-DNA lines used to test genes identified in a genome wide association studies (GWAS) were obtained from the Arabidopsis biological resource center and confirmed homozygous for the insertion by PCR ^42^.

### Plant Growth Conditions

For experimental plants, seeds collected from the bulking were stratified for 5 days as described above and planted in 2 ¼ inch pots filled with a 1:2 sand (Quikrete), BX potting mix (Pro-mix) mix with thin layer of 100% BX potting mix on top. The 60/30 mix was used for faster drainage so that drought conditions were met quickly when water was withheld, while the layer of 100% BX was used to aid in seed germination and image analysis. Potting substrate was supplemented with 2x miracle grow and marathon granules for added nutrients and pest management. Plants were grown in a Conviron chamber with environmental conditions as described above. Plants were well-watered for 10 days before being moved to a cold room at 4°C for a 5-week vernalization. Plants used in experiments for validating GWAS results did not undergo a vernalization treatment and lacked the layer of 100% BX.

### Drought treatments

Well-watered plants were bottom watered twice daily, 12 hours apart, for 10 mins using an automated watering system comprised of flood trays, water reservoirs, and water pumps on timers. Plants in the juvenile drought treatment experienced a 10-day drought starting 47 days after planting (2 days after being moved back to growth conditions following vernalization). During the drought, plants were bottom watered for 10 mins every 84 hours. The timing of adult drought treatments was determined based on flowering time during the bulking for each genotype to ensure plants began drought prior to flowering but after transitioning to the adult vegetative phase. Accessions were binned into four groups of 40 based on flowering time, accessions in group 1 began adult drought treatments 55 days after planting, group 2, 60 days after planting and groups 3 and 4, 65 days after planting (Table S1). miR156OE and MIM156 lines were put in the same group as the wild-type Col-0. Adult drought treatments were conducted the same way as juvenile drought treatments for 10-days with watering intervals of 84 hours. Following drought treatments, all plants returned to well-watered conditions with a watering interval of 12 hours.

Plants used in experiments for validating GWAS results experiments were well-watered by bottom watering for 10 mins once a day. During drought treatments, water was withheld for 10 days. Juvenile drought began when the first two true leaves were visible, and adult drought began when leaf 5 was visible. Following drought treatments, all plants returned to well-watered conditions with a watering interval of once per day. Volumetric water content of growth media was measured hourly with METER Group EC-5 sensors (Fig S1).

### Phenotyping for the timing of vegetative phase change

We used the first leaf with abaxial trichomes as a marker for the onset of vegetative phase change ^43,44^. A minimum of 3 replicates per genotype were measured for all natural accessions in the well-watered conditions and a subset of 30 genotypes were measured in the juvenile drought treatment to look at plasticity in the timing of VPC. Daily images taken of the rosettes by RaspberryPis were used to convert the timing of VPC from leaf number to days by noting the date when the first leaf with abaxial trichomes was visible.

### Phenotyping for rosette growth

To study rosette growth responses to drought we measured rosette diameter on the first and last day of drought treatments, and the corresponding days for well-watered control plants. Measurements were taken with a combination of PlantCV ^45^ using the “*Arabidopsis* multi-plant workflow” and threshold settings between 110-120 and manual measurements in FIJI ^46^. Change in size during drought was calculated by subtracting the starting diameter from the final diameter.

### Phenotyping for fruit production

Plants were harvested when the first fruit became mature determined by a yellowing in color as Boyes *et al.*, 2001 determined that flowering is complete around the same day the first silique shatters in Col-0. Shoots were cut at the soil surface and Inflorescences and rosettes were separated and laid flat on black fabric and imaged with a Nikon D3200. Images were analyzed for estimating fruit count following Vasseur et al. 2018 ^47^. We found the Vasseur model predicted fruit count of natural accessions from images in our experiment well, with an R^2^ of 0.91 for the linear regression model (manual ∼ predicted) between predicted versus manually counted fruits. For mutant genotypes, we calculated a new model based on manually counted samples from the mutants (nbBranches x 0.040261074 + EndPointsVox x 0.134424186 + SlabVox x 0.004873814) which had an R^2^ of 0.94 for the linear regression model (manual ∼ predicted) compared to an R^2^ of 0.78 when using the Vasseur model for this population.

### Isotope discrimination analysis

Plants for isotope analysis were grown as previously described. The two newest fully expanded leaves for three replicates of each genotype were sampled at the end of the juvenile drought treatment, the end of the adult drought treatment, and at corresponding time points for well-watered controls. Leaf tissue was dried at 60°C for a minimum of 48 hours, then pooled by genotype and treatment. Samples were ground to a fine homogenous powder using a Benchmark Bead Bug 6 Microtube Homogenizer with a 1.6 mm stainless steel ball. Between 0.09 and 3.1 mg (typically ∼2mg per sample) were weighed into 8 x 5 mm tin capsules using a Sartorius MCE3.6P microbalance. Carbon and nitrogen isotope ratios and percent composition were measured by the Cornell University Stable Isotope Lab using a Thermo Delta V isotope ratio mass spectrometer (IRMS) interfaced to a NC2500 elemental analyzer. The primary reference scale used for δN was atmospheric air and for δC was Vienna Pee Dee Belemnite.

### miR156 mutant analysis

To assess the role of miR156 expression in phase-specific responses to drought in fruit production we calculated the percent of control fruits in each drought treatment for Col-0 wildtype, 35S:miR156 OE and MIM156 lines as 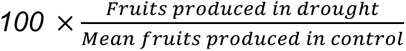 to account for variation in fruit production among mutants in well-watered control conditions. We used two-way ANOVA with Genotype, Treatment and their interaction as fixed effects. When a significant effect was detected, pairwise comparisons of estimated marginal means (EMMs) were conducted using Tukey’s HSD adjustment for multiple comparisons using the emmeans package in R.

### Principle component analysis

Principal component analysis (PCA) was used to summarize phenotypic variation across genotypes and treatments. The traits included in the analysis were genotypic means of fruits produced, rosette diameter growth, δ^13^C, timing of VPC based on first leaf with abaxial trichomes, and flowering time in well-watered control, juvenile drought and adult drought conditions. Genotypes with missing data were removed from the analysis and variables were centered and scaled to unit variance prior to PCA. PCA was performed using the prcomp() function in R and the first two principal components were visualized to assess patterns of variation among genotypes.

### Bioclimatic associations with the timing of VPC

Bioclimatic variables corresponding to the latitude and longitude of each accession’s origin were obtained from CHELSEA-daily version 2.1 ^48^. We used linear regression to look for correlations between the bioclimatic variables and the timing of VPC. To look at seasonally relevant climate traits when *A. thaliana* is growing, we calculated mean precipitation, temperature and vapor pressure deficit from monthly values across September, October and November which we call fall and February, March and April which we refer to as spring. We further used mixed models controlling for genetic similarity to confirm these relationships were not due to kinship as described in Gamba *et al*. 2024^31^. Analyses were conducted both globally, across all accessions, and within regions. Given the wide variation in both climate and VPC timing among accessions from the Iberian Peninsula (Spain and Portugal) and high number of accessions from Germany, and the fact that clines often change between parts of the *Arabidopsis* range ^31^. We performed additional analyses on these subsets to explore region-specific patterns.

### Genome wide association analysis

Genome wide association studies (GWAS) were conducted using the GWA-Portal tool (Seren 2018) with the Accelerated Mixed Model (AMM) protocol and 1001 “Full sequence” dataset (TAIR9) ^49^. GWAS was performed separately for each trait, using mean trait values per genotype. The datasets included 158 *Arabidopsis* accessions with genotype and phenotype data. A minor allele frequency (MAF) threshold of 0.05 was applied. Candidate genes near the top SNPs were identified using Araport11 annotations and the TAIR Gene Search tool (arabidopsis.org) examining a 10 kb window around each SNP.

### Candidate gene follow-up using T-DNA insertion lines and natural variants

48 T-DNA insertion lines representing mutations in 26 candidate genes (Table S2) identified from GWAS were selected using the Salk Institute Genomic Analysis Laboratory (SIGnAL) T-DNA Express tool and seeds obtained from the Arabidopsis Biological Resources Center. Some candidate genes identified in early GWAS results were no longer near top SNPs of current iterations, but were still included in this analysis. Plants were grown together in a Conviron growth chamber in conditions described above for genotyping and bulking. Individual plants were genotyped by PCR using the Thermo Fisher Phire Plant Direct PCR Kit and primers designed with the SIGnAL T-DNA Primer Design tool. Seeds were collected from the plants confirmed homozygous for the T-DNA insertion (21 lines) and used in subsequent experiments. In addition, we used the Polymorph 1001 tool (1001genomes.org) to identify accessions in our study with natural loss of function (LOF) alleles to compare phenotypes of accessions with the Col-0 reference allele and with putative LOF alleles at candidate genes.

### Statistical Analyses

All statistical analyses were performed in R version 4.2.3 ^50^. Figures were generated using the ggplot2 package in R. Phenotypic differences across control, juvenile and adult drought treatments were assessed using two-way analysis of variance (ANOVA) with genotype, environment, and their interaction as fixed effects, followed by Tukey’s post hoc test. Comparisons of isotope discrimination between drought treated and corresponding well-watered control plants were conducted using Student’s T-test. Correlations between continuous variables were evaluated using linear regression models (R function lm). To test for significant differences between mutant lines and the wild-type control, we used EMMs calculated using the emmeans package. Dunnett-adjusted post hoc contrasts were then used to compare each mutant genotype to the WT control within each treatment condition.

### Tests of selection on top GWAS SNPs

If genetic loci are under spatially varying selection, we expect they will exhibit greater spatial variation in allele frequency than the genomic background. By contrast, loci under balancing or contributing to traits under consistent stabilizing selection may show less spatial variation than the genomic background. We tested for this pattern across a range of spatial scales using published calculations of scale-specific genetic variance across the native accessions in the 1001 Genomes panel ^34^. We took either the top 100 or 1,000 SNPs from GWAS for timing of VPC and fitness in each drought treatment and asked if the mean variance at a range of spatial scales was greater or less than expected from 100 permutations of SNP labels around the genome.

### Natural climate relationships during germination, early vegetative development and reproduction

To compare climate conditions during key developmental phases, we used location and timing data for natural populations from Norwich, Cologne, Halle and Valencia as reported in Wilczek *et al.* 2010^40^. Daily maximum and minimum temperature and precipitation data for each site were obtained using the nasapower package in R. For each population and cohort, we summarized climate conditions over three developmental windows: the two weeks preceding germination, two weeks following germination (representing early vegetative development) and the two weeks following bolting (representing the reproductive phase). Mean temperature was calculated as the average of daily maximum and minimum temperatures over each period. Linear regression was used to assess consistency in environmental variables across developmental phases, sites and cohorts.

## Supplemental Figures

**Figure S1.**
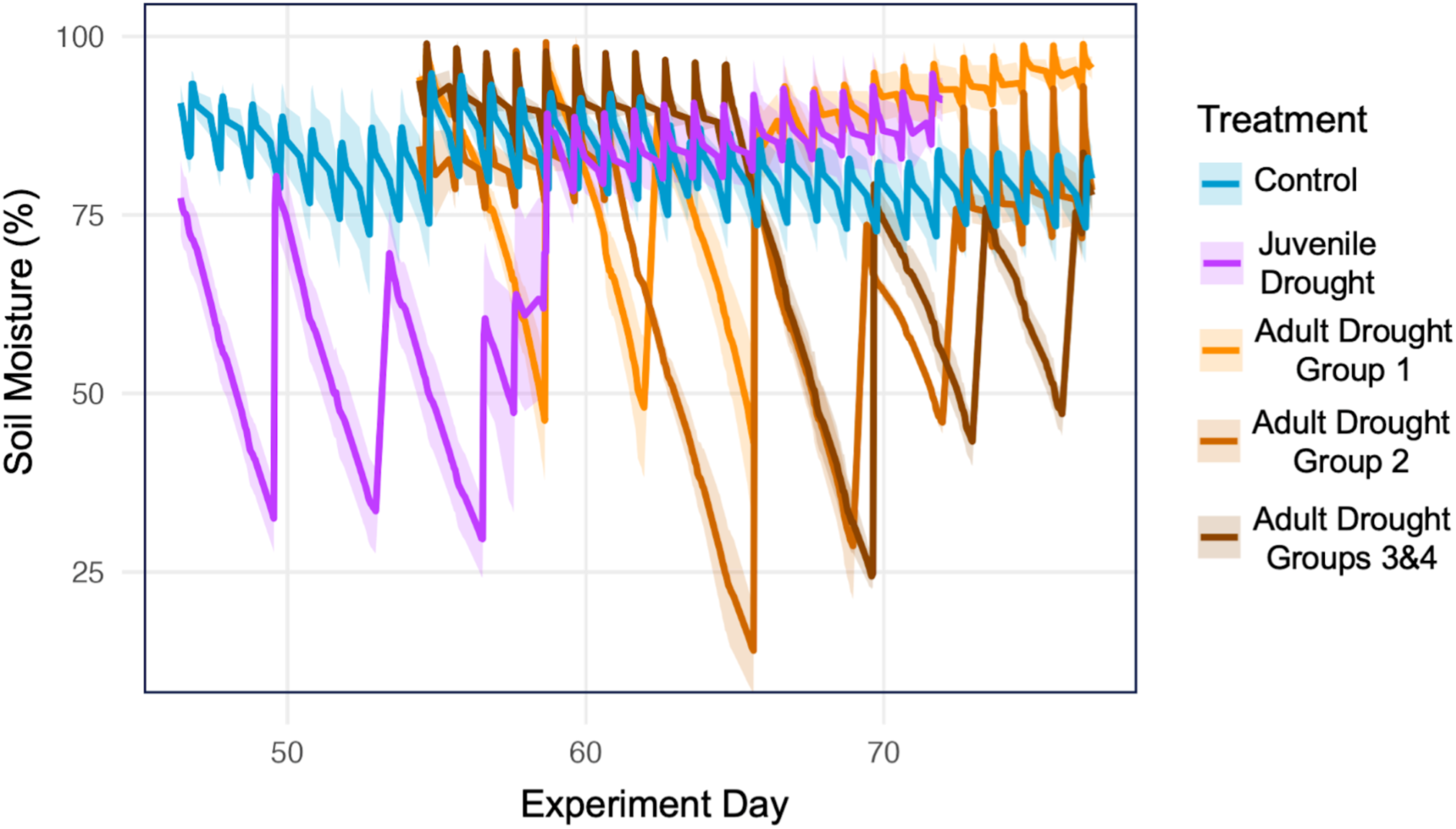
Soil water content during drought periods of the experiment for each treatment group. Solid lines show the mean percent of maximum soil capacity every hour measured by 2-4 soil moisture meters with shading showing the standard error. Blue shows well-watered control treatment, purple shows the juvenile drought treatment where drought begins on day 46, light orange shows the adult drought treatment for genotypes in group 1 where drought begins on day 55, medium orange shows adult drought group 2 where drought begins on day 60 and brown shows adult drought for groups 3 and 4 where drought begins on day 65.

**Figure S2.**
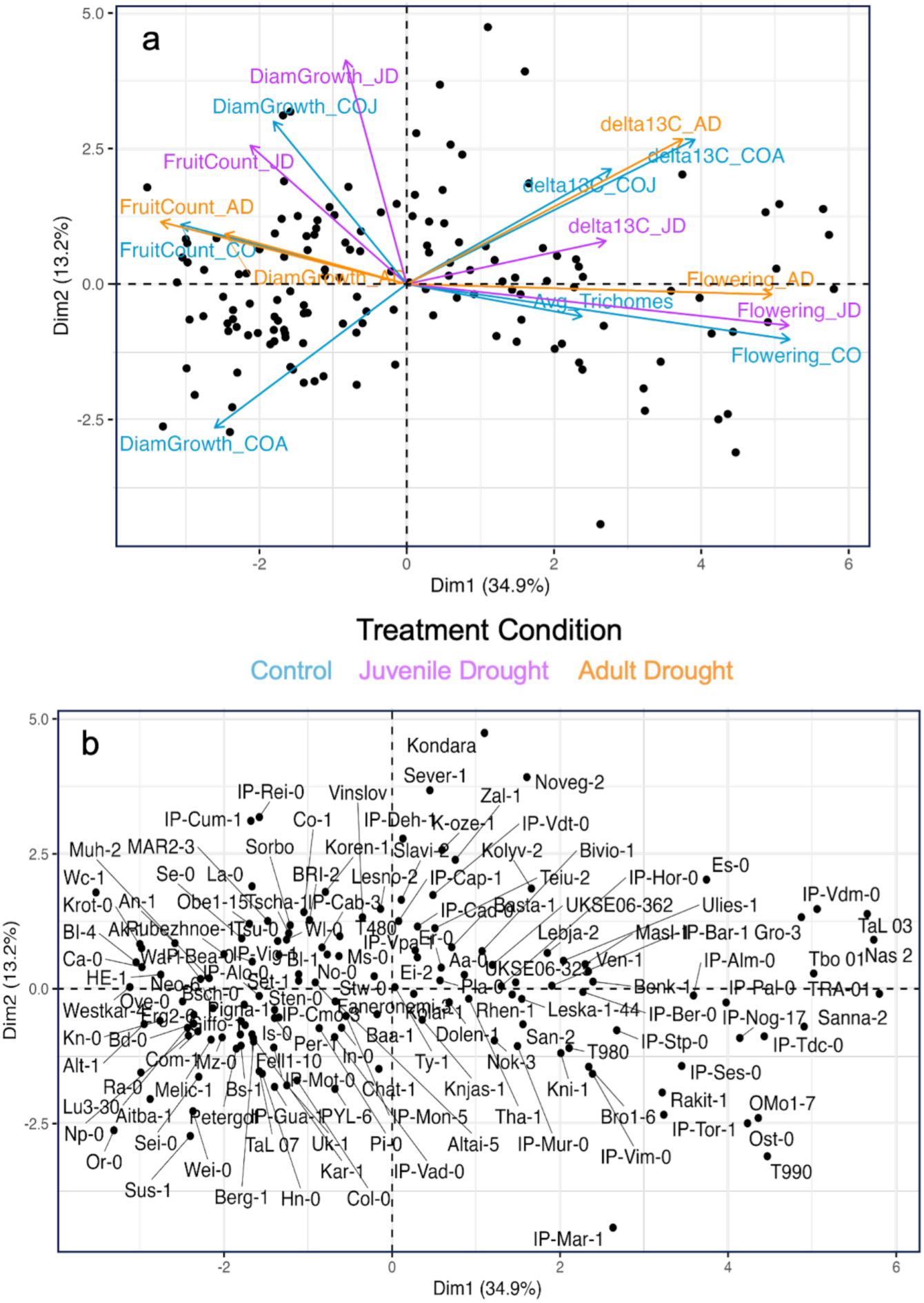
Principal component analysis (PCA) of growth, physiology, fitness and developmental phenotypes for *A. thaliana* accessions subjected to drought during the juvenile vegetative or adult vegetative phase and well-watered controls. (a-b) Biplot of PC 1 and 2 which describe 34.9% and 13.2% of phenotypic variation respectively. Black dots represent individual accessions (a-b) and arrows (a) represent trait loadings with phenotypes from well-watered control (CO) conditions in blue, juvenile drought (JD) in purple and adult drought (AD) in orange. For control traits measured at a specific developmental stage, COJ indicates control juvenile and COA, control adult.

**Figure S3.**
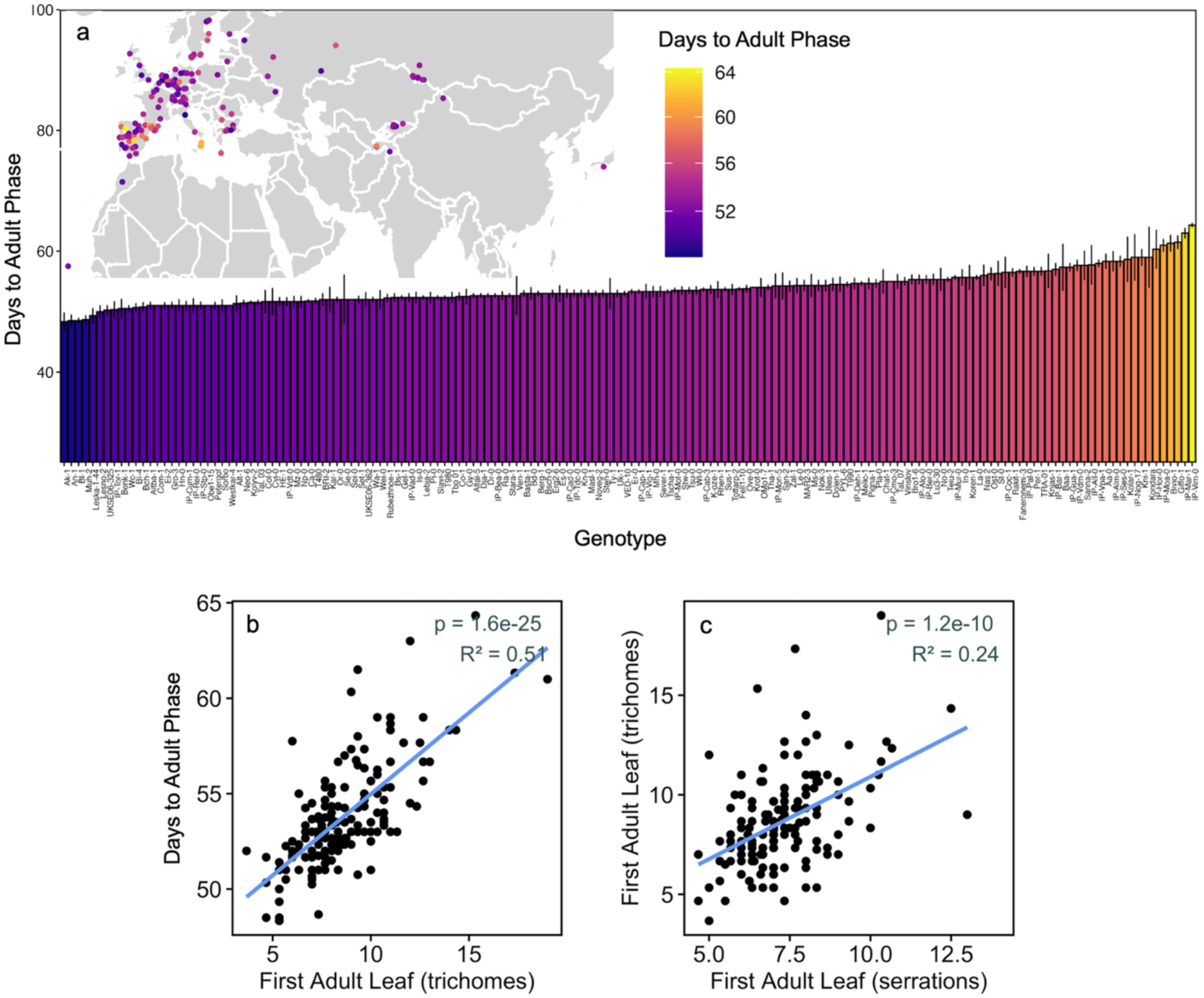
Natural variation in the timing of vegetative phase change (VPC). Timing of VPC based on the days between planting and when the first adult leaf with abaxial trichomes was present for 158 natural accessions used in this study (A). Accessions are plotted on the map in the location where they originated from, and points are colored based on days until the first adult leaf is visible. Bars represent the mean ± SE for a minimum of 3 replicates per genotype. Relationships between the timing of VPC measured as days till the first adult leaf is visible, first adult leaf based on abaxial trichomes and based on serrations (B, C). Results from linear regression analysis displayed in the top right corner of each plot.

**Figure S4.**
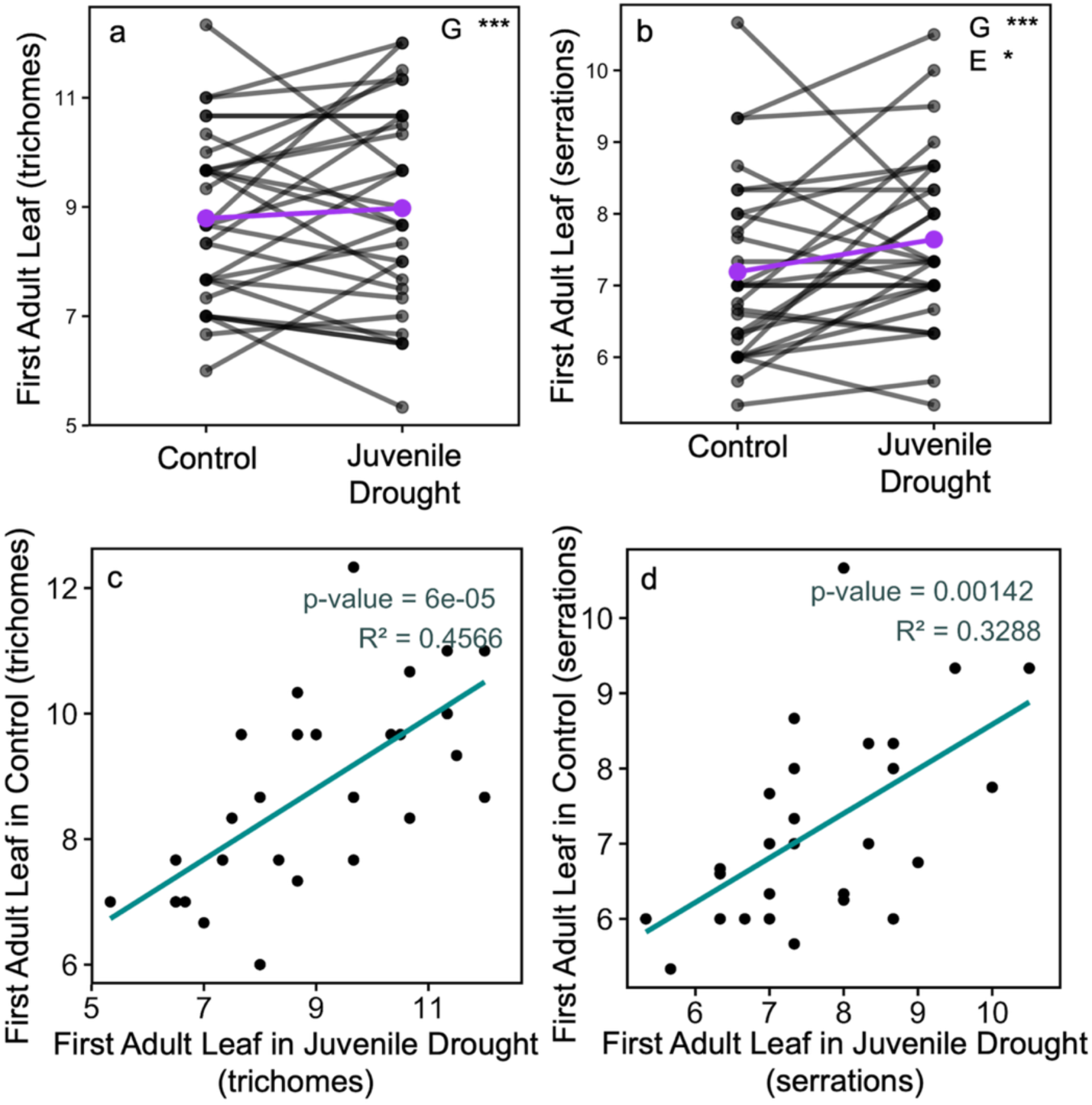
Plasticity in the timing of vegetative phase change between well-watered control and juvenile drought treatments. Black circles represent the mean first adult leaf based on abaxial trichomes (a,c)) or serrations (b,d) for each of the 30 accessions examined in both treatments. Black lines (a-b) connect each genotype between treatments and purple circles and lines represent the mean for all genotypes. ANOVA results depicted in the top right corner of each plot with *** and * indicating *p <* 0.001 and 0.05 respectively. (c-d) Teal lines represent the line of best fit and results from linear regression analysis are displayed in the top right corner of each plot.

**Figure S5.**
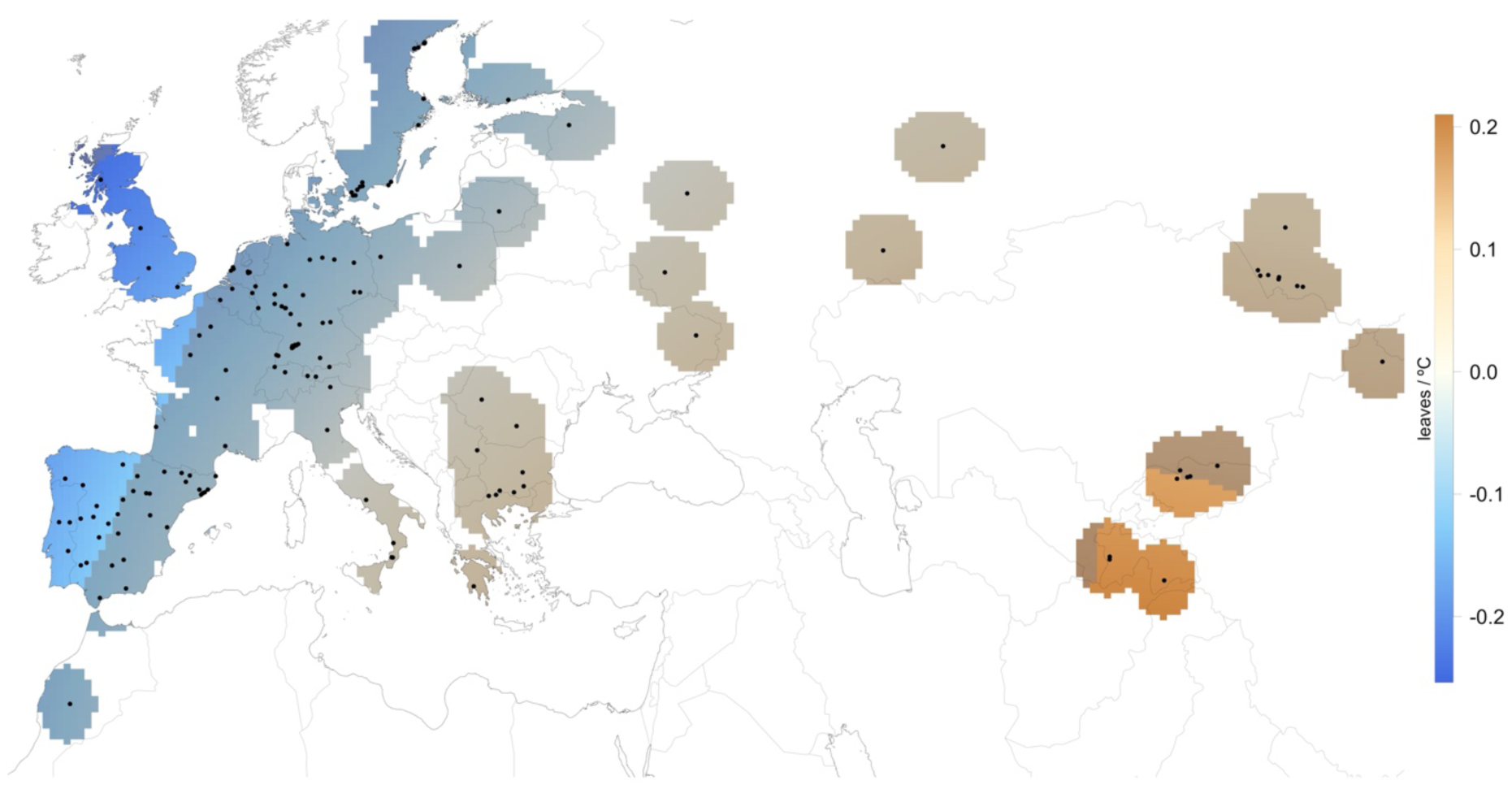
Results of generalized-additive-model (GAM) to detect significant trends between the timing of VPC and mean spring temp

**Figure S6.**
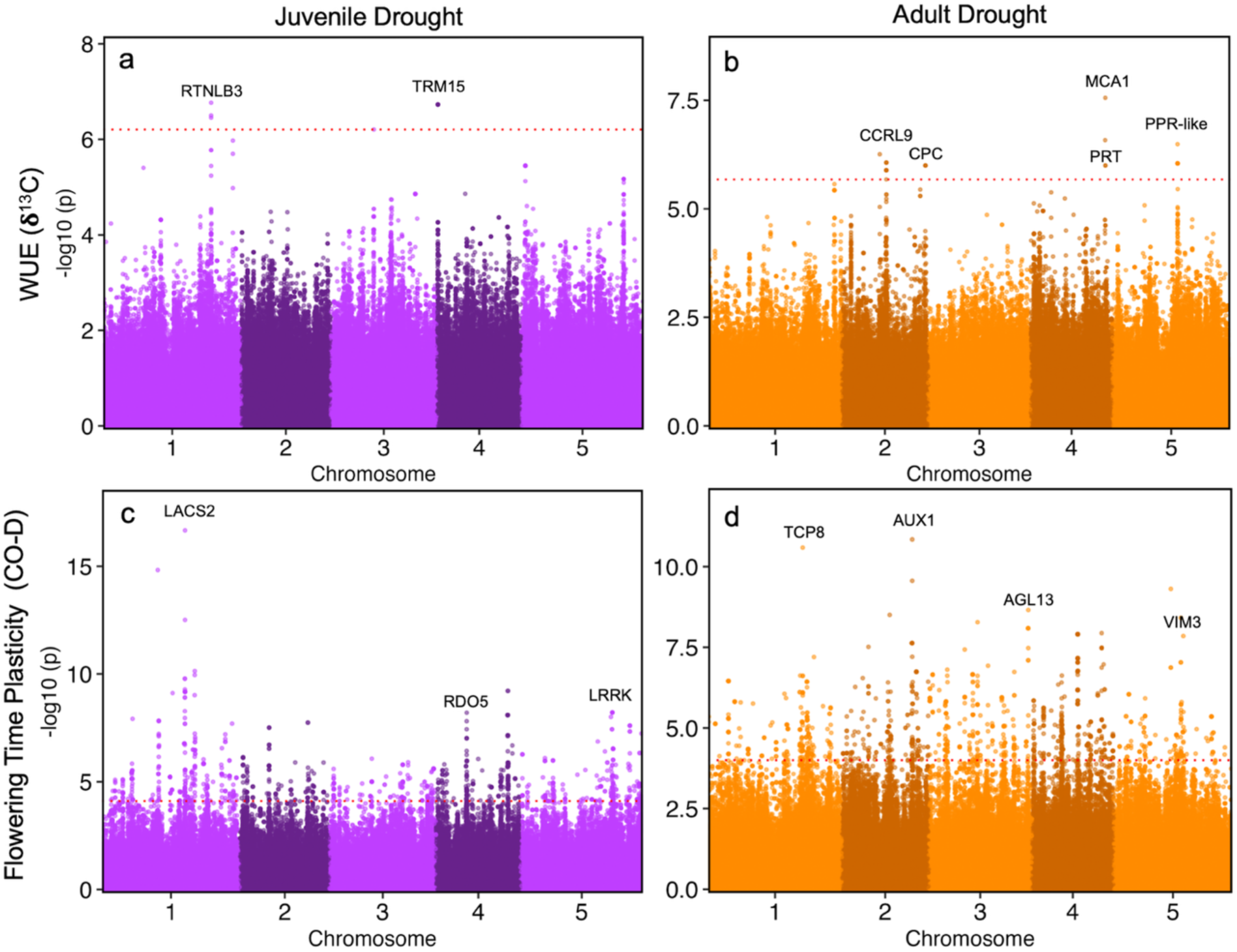
Manhattan plots of genome wide association (GWA) analysis on fitness responses to drought during juvenile (a, c) and adult (b, d) phases of development. Manhattan plots show the association of each SNP with water use efficiency (WUE) (a, b) or flowering time plasticity between well-watered control and drought treatments (c, d). Red dotted lines indicate significance threshold based on q-values with Benjamin Hochberg FDR correction with *p* < 0.1. Annotations indicate candidate genes associated with top SNPs.

**Figure S7.**
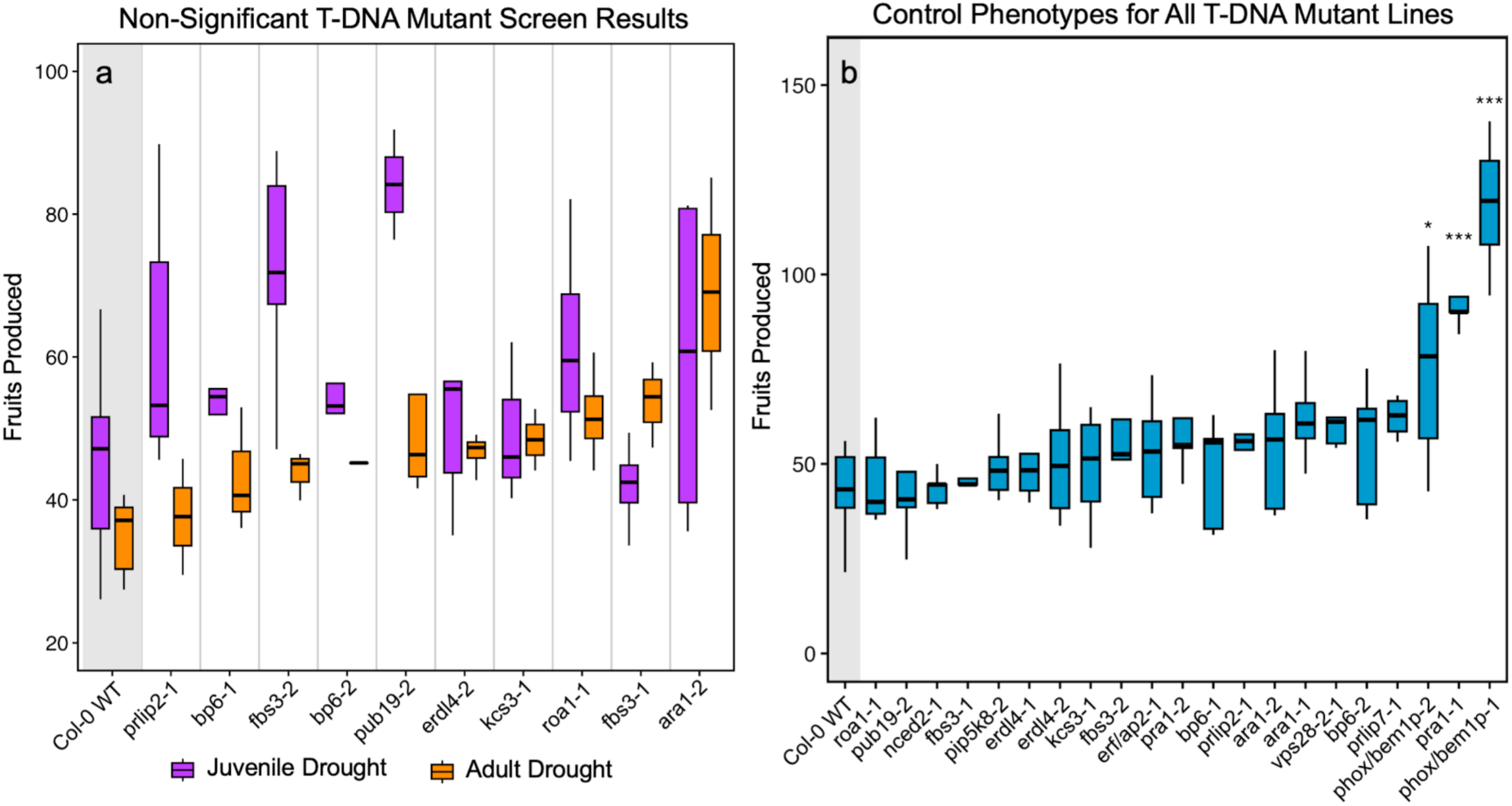
Fruits produced by Col-0 wild-type and T-DNA insertion lines with no significant differences from Col-0 based on Dunnett-adjusted contrasts when exposed to drought during the juvenile (purple) or adult (orange) phase (a). Fruits produced by Col-0 and all T-DNA lines tested for phase-specific drought responses in well-watered control conditions (b). Results from Dunnett-adjusted contrasts between Col-0 and each T-DNA line in each condition indicated above each boxplot where ***, **, and * indicate *p <* 0.001, 0.01 and 0.05 respectively.

**Figure S8.**
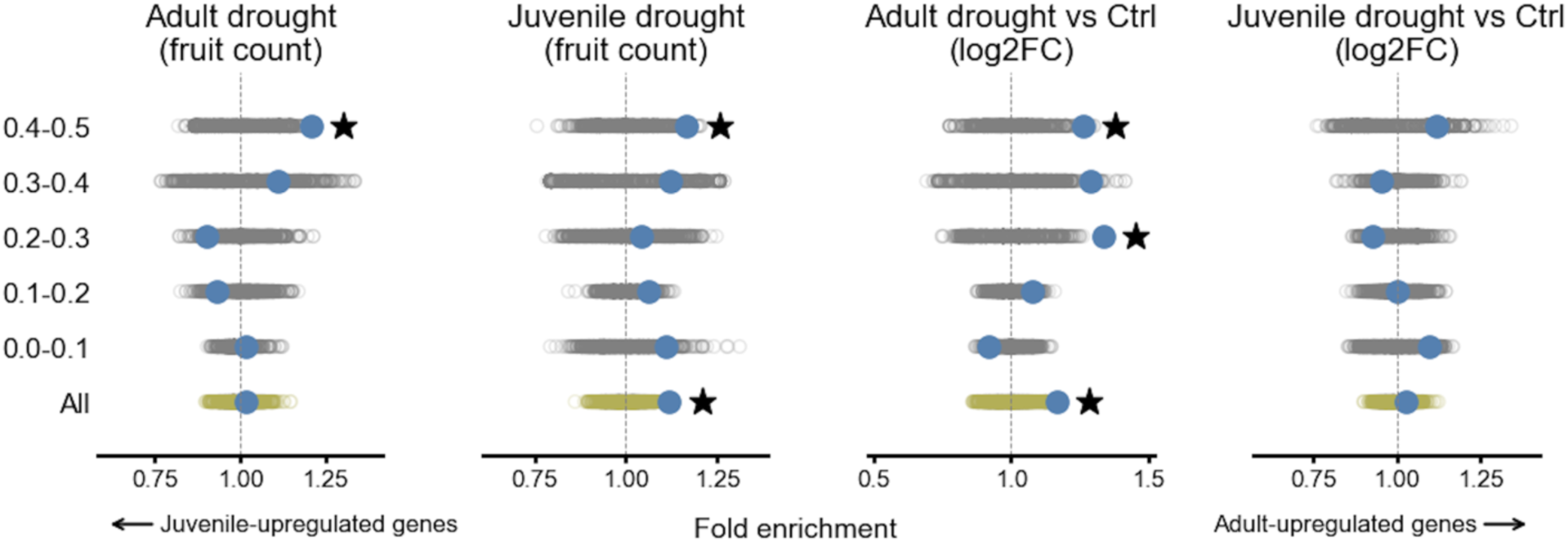
Juvenile and adult expression enrichment of drought related GWAS SNPs

**Figure S9.**
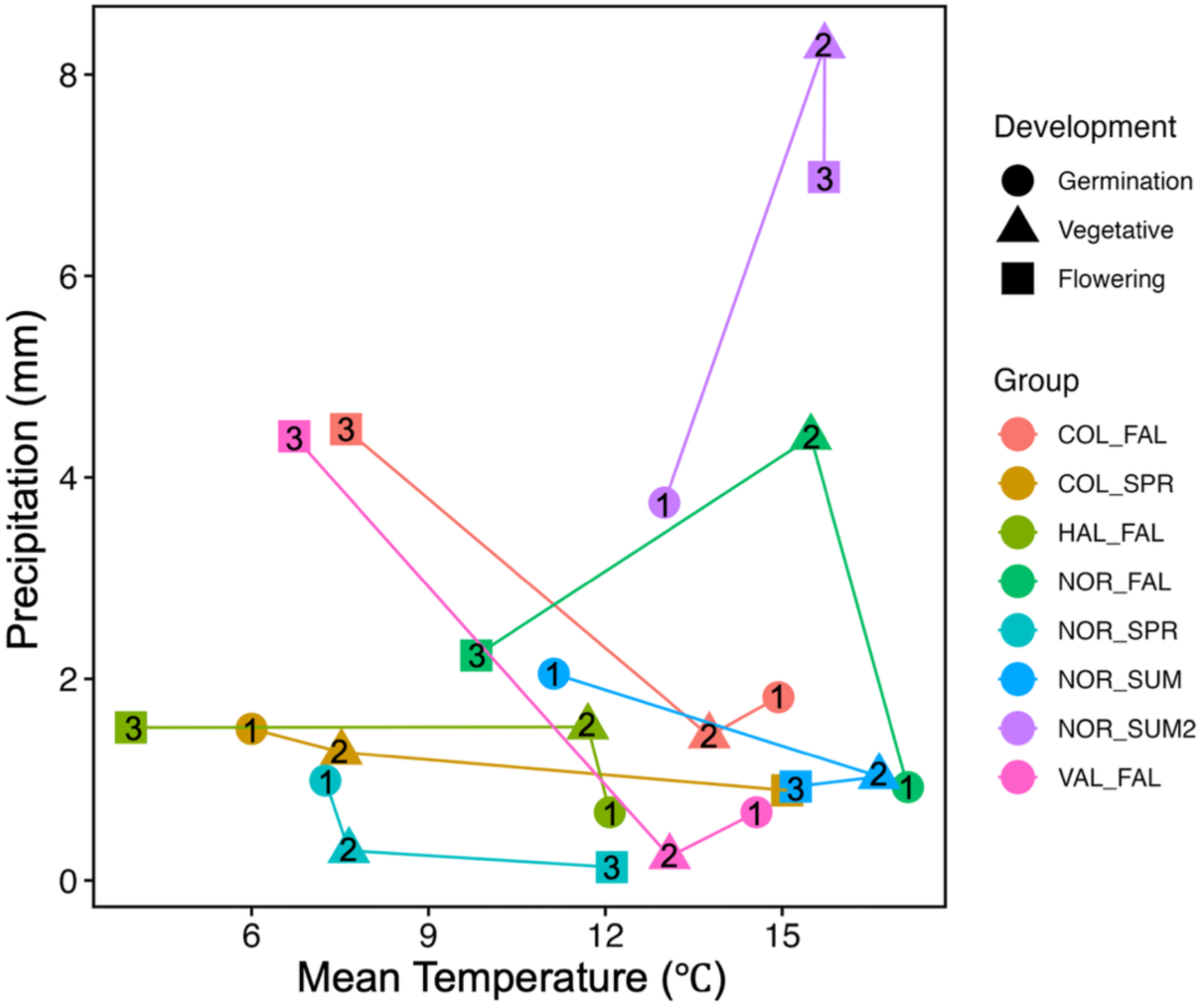
Precipitation and temperature conditions for natural *A. thaliana* populations from Norwich, Cologne, Halle and Valencia reported in Wilczek *et al.* 2010 during different phases of development. Circles labeled “1” represent conditions around germination, triangles labeled “2” represent conditions during the early vegetative phase and squares labeled “3” represent conditions during flowering. Conditions for each cohort of plants (grouped by location and season) are shown in different colors with lines connecting the points for each developmental phase. Linear regression was used to assess consistency in environmental variables across developmental phases, sites and cohorts. Mean temperature during germination was significantly correlated with early vegetative temperature (*p <* 0.05, R^2^ = 0.6) but not with temperature during flowering (*p =* 0.24). Similarly, precipitation during germination was significantly correlated with early vegetative conditions (*p <* 0.05, R^2^ = 0.55) but not with precipitation during flowering (*p =* 0.11) indicating that conditions during germination are predictive of conditions during early vegetative development, but not reproduction.

## References

1. Lawrence-Paul, E. & Lasky, J. Ontogenetic changes in ecophysiology are an understudied yet important component of plant adaptation. Am. J. Bot. 111, e16294 (2024).

2. Donohue, K. Germination timing influences natural selection on life-history characters in arabidopsis thaliana. Ecology 83, 1006–1016 (2002).

3. Stinchcombe, J. R. et al. A latitudinal cline in flowering time in Arabidopsis thaliana modulated by the flowering time gene FRIGIDA. Proc. Natl. Acad. Sci. 101, 4712–4717 (2004).

4. Korves, T. M. et al. Fitness effects associated with the major flowering time gene FRIGIDA in Arabidopsis thaliana in the field. Am. Nat. 169, (2007).

5. Kenney, A. M., McKay, J. K., Richards, J. H. & Juenger, T. E. Direct and indirect selection on flowering time, water-use efficiency (WUE, δ13C), and WUE plasticity to drought in *Arabidopsis thaliana*. Ecol. Evol. 4, 4505–4521 (2014).

6. Burghardt, L. T. et al. Modeling the Influence of Genetic and Environmental Variation on the Expression of Plant Life Cycles across Landscapes. Am Nat 185, 212–227 (2015).

7. Austen, E. J., Rowe, L., Stinchcombe, J. R. & Forrest, J. R. K. Explaining the apparent paradox of persistent selection for early flowering. New Phytol. 215, 929–934 (2017).

8. Dayrell, R. L. C. et al. Ontogenetic shifts in plant ecological strategies. Funct. Ecol. 32, 2730–2741 (2018).

9. Bond, W. J., Lee, W. G. & Craine, J. M. Plant structural defences against browsing birds: a legacy of New Zealand’s extinct moas. Oikos 104, 500–508 (2004).

10. Howard, J., Cameron, E., Bellve, A., Baba, Y. & Wright, S. New Zealand divaricate plant species: Tensile strength and Remote Island occurrence. Austral Ecol. 47, 1091–1100 (2022).

11. Stearns, S. C. Trade-Offs in Life-History Evolution. Funct. Ecol. 3, 259–268 (1989).

12. Mojica, J. P., Lee, Y. W., Willis, J. H. & Kelly, J. K. Spatially and temporally varying selection on intrapopulation quantitative trait loci for a life history trade-off in Mimulus guttatus. Mol. Ecol. 21, 3718–3728 (2012).

13. Bazzaz, F. A., Chiariello, N. R., Coley, P. D. & Pitelka, L. F. Allocating Resources to Reproduction and Defense. BioScience 37, 58–67 (1987).

14. Young, T. P. & Augspurger, C. K. Ecology and evolution of long-lived semelparous plants. Trends Ecol. Evol. 6, 285–289 (1991).

15. Lundgren, M. R. & Des Marais, D. L. Life History Variation as a Model for Understanding Trade-Offs in Plant–Environment Interactions. Curr. Biol. 30, R180–R189 (2020).

16. Wu, G. & Poethig, R. S. Temporal regulation of shoot development in *Arabidopsis thaliana* by miR156 and its target SPL3. Development 133, 3539–3547 (2006).

17. Willmann, M. R. & Poethig, R. S. Conservation and evolution of miRNA regulatory programs in plant development. Curr. Opin. Plant Biol. 10, 503–511 (2007).

18. Wu, G. et al. The sequential action of miR156 and miR172 regulates developmental timing in *Arabidopsis*. Cell 138, 750–759 (2009).

19. Hudson, C. J. et al. Genetic Control of Heterochrony in Eucalyptus globulus. G3 GenesGenomesGenetics 4, 1235–1245 (2014).

20. Xu, M. et al. Developmental functions of miR156-regulated SQUAMOSA PROMOTER BINDING PROTEIN-LIKE (SPL) genes in *Arabidopsis thaliana*. PLOS Genet. 12, e1006263 (2016).

21. Lucani, C. J., Brodribb, T. J., Jordan, G. J. & Mitchell, P. J. Juvenile and adult leaves of heteroblastic Eucalyptus globulus vary in xylem vulnerability. Trees 33, 1167–1178 (2019).

22. Visentin, I. et al. A novel strigolactone-miR156 module controls stomatal behaviour during drought recovery. Plant Cell Environ. 43vis, 1613–1624 (2020).

23. Lawrence, E. H., Springer, C. J., Helliker, B. R. & Poethig, R. S. MicroRNA156-mediated changes in leaf composition lead to altered photosynthetic traits during vegetative phase change. New Phytol. 231, 1008–1022 (2021).

24. Lawrence, E. H., Springer, C. J., Helliker, B. R. & Poethig, R. S. The carbon economics of vegetative phase change. Plant Cell Environ. 45, 1286–1297 (2022).

25. Lawrence-Paul, E. H., Poethig, R. S. & Lasky, J. R. Vegetative phase change causes age-dependent changes in phenotypic plasticity. New Phytol. 240, 613–625 (2023).

26. Mao, H.-D. et al. Genome-wide analysis of the SPL family transcription factors and their responses to abiotic stresses in maize. Plant Gene 6, 1–12 (2016).

27. Zhao, Y. et al. The SPL transcription factor TaSPL6 negatively regulates drought stress response in wheat. Plant Physiol. Biochem. 206, 108264 (2024).

28. Lasky, J. R. et al. Natural Variation in Abiotic Stress Responsive Gene Expression and Local Adaptation to Climate in Arabidopsis thaliana. Mol. Biol. Evol. 31, 2283–2296 (2014).

29. Lee, G., et al. A large-effect fitness trade-off across environments is explained by a single mutation affecting cold acclimation. Proc. Natl. Acad. Sci. 121, e2317461121 (2024).

30. He, J. et al. Threshold-dependent repression of SPL gene expression by miR156/miR157 controls vegetative phase change in *Arabidopsis thaliana*. PLOS Genet. 14, e1007337 (2018).

31. Gamba, D. et al. The genomics and physiology of abiotic stressors associated with global elevational gradients in Arabidopsis thaliana. New Phytol. 244, 2062–2077 (2024).

32. Zhan, X. et al. An Arabidopsis PWI and RRM motif-containing protein is critical for pre-mRNA splicing and ABA responses. Nat. Commun. 6, 8139 (2015).

33. Hu, L. et al. Distinct function of SPL genes in age-related resistance in Arabidopsis. PLOS Pathog. 19, e1011218 (2023).

34. Lasky, J. R., Takou, M., Gamba, D. & Keitt, T. H. Estimating scale-specific and localized spatial patterns in allele frequency. Genetics 227, iyae082 (2024).

35. Axtell, M. J. & Bowman, J. L. Evolution of plant microRNAs and their targets. Trends Plant Sci. 13, 343–349 (2008).

36. Bahuguna, R. N. et al. Physiological and biochemical characterization of NERICA-L-44: a novel source of heat tolerance at the vegetative and reproductive stages in rice. Physiol. Plant. 154, 543–559 (2015).

37. Driedonks, N., Rieu, I. & Vriezen, W. H. Breeding for plant heat tolerance at vegetative and reproductive stages. Plant Reprod. 29, 67–79 (2016).

38. Varoquaux, N. et al. Transcriptomic analysis of field-droughted sorghum from seedling to maturity reveals biotic and metabolic responses. Proc. Natl. Acad. Sci. U. S. A. 116, 27124– 27132 (2019).

39. Zhao, J., Doody, E. & Poethig, R. S. Reproductive competence is regulated independently of vegetative phase change in Arabidopsis thaliana. Curr. Biol. 33, 487–497 (2023).

40. Wilczek, A. M. et al. Genetic and physiological bases for phenological responses to current and predicted climates. Philos. Trans. R. Soc. B Biol. Sci. 365, 3129–3147 (2010).

41. Franco-Zorrilla, J. M. et al. Target mimicry provides a new mechanism for regulation of microRNA activity. Nat. Genet. 39, 1033–1037 (2007).

42. Krysan, P. J., Young, J. C. & Sussman, M. R. T-DNA as an Insertional Mutagen in Arabidopsis. Plant Cell 11, 2283–2290 (1999).

43. Telfer, A., Bollman, K. M. & Poethig, R. S. Phase change and the regulation of trichome distribution in *Arabidopsis thaliana*. Development 124, 645–654 (1997).

44. Willmann, M. R. & Poethig, R. S. The effect of the floral repressor FLC on the timing and progression of vegetative phase change in Arabidopsis. Development 138, 677–685 (2011).

45. Gehan, M. A. et al. PlantCV v2: Image analysis software for high-throughput plant phenotyping. PeerJ 5, e4088 (2017).

46. Schindelin, J., et al. Fiji: An open-source platform for biological-image analysis. Nat. Methods 9, 676–682 (2012).

47. Vasseur, F. et al. Climate as a driver of adaptive variations in ecological strategies in Arabidopsis thaliana. Ann. Bot. 122, 935–945 (2018).

48. Karger, D. N. et al. Climatologies at high resolution for the earth’s land surface areas. Sci. Data 4, 170122 (2017).

49. Alonso-Blanco, C. et al. 1,135 Genomes Reveal the Global Pattern of Polymorphism in Arabidopsis thaliana. Cell 166, 481–491 (2016).

50. R Core Team. R: A Language and Environment for Statistical Computing. R Foundation for Statistical Computing (2021).

